# Population analysis of *Legionella pneumophila* reveals the basis of resistance to complement-mediated killing

**DOI:** 10.1101/2020.08.14.250670

**Authors:** Bryan A. Wee, Joana Alves, Diane S. J. Lindsay, Ross L. Cameron, Amy Pickering, Jamie Gorzynski, Jukka Corander, Pekka Marttinen, Andrew J. Smith, J. Ross Fitzgerald

**Author notes:** These authors contributed equally (Bryan A. Wee, Joana Alves).

## Abstract

*Legionella pneumophila* is the most common cause of the severe respiratory infection known as Legionnaires’ disease. *L. pneumophila* is typically a symbiont of free-living amoeba, and our understanding of the bacterial factors that determine human pathogenicity is limited. Here we carried out a population genomic study of 900 *L. pneumophila* isolates from human clinical and environmental samples to examine their genetic diversity, global distribution and the basis for human pathogenicity. We found that although some clones are more commonly associated with clinical infections, the capacity for human disease is representative of the breadth of species diversity. To investigate the bacterial genetic basis for human disease potential, we carried out a genome-wide association study that identified a single gene (*lag-1*), to be most strongly associated with clinical isolates. Molecular evolutionary analysis showed that *lag-1*, which encodes an *O*-acetyltransferase responsible for lipopolysaccharide modification, has been distributed horizontally across all major phylogenetic clades of *L. pneumophila* by frequent recent recombination events. Functional analysis revealed a correlation between the presence of a functional *lag-1* gene and resistance to killing in human serum and bovine broncho-alveolar lavage. In addition, *L. pneumophila* strains that express *lag-1* escaped complement-mediated phagocytosis by neutrophils. Importantly, we discovered that the expression of *lag-1* confers the capacity to evade complement-mediated killing by inhibiting deposition of classical pathway molecules on the bacterial surface. In summary, our combined population and functional analyses identified *L. pneumophila* genetic traits linked to human disease and revealed the molecular basis for resistance to complement-mediated killing, a previously elusive trait of direct relevance to human disease pathogenicity.

**Significance:** *Legionella pneumophila* is an environmental bacterium associated with a severe pneumonia known as Legionnaires’ disease. A small number of *L. pneumophila* clones are responsible for a large proportion of human infections suggesting they have enhanced pathogenicity. Here, we employed a large-scale population analysis to investigate the evolution of human pathogenicity and identified a single gene (*lag-1*) that was more frequently found in clinical isolates. Functional analysis revealed that the *lag-1*-encoded *O*-acetyltransferase, involved in modification of lipopolysaccharide, conferred resistance to the classical pathway of complement in human serum. These findings solve a long-standing mystery in the field regarding *L. pneumophila* resistance to serum killing, revealing a novel mechanism by which *L. pneumophila* may avoid immune defences during infection.

## Introduction

*Legionella pneumophila* is a γ-proteobacterial species that parasitises free-living amoeba in freshwater environments (Hilbi *et al*., 2011). *L. pneumophila* hijacks the phagocytic process in amoebae and human alveolar macrophages by subverting host cellular mechanisms to promote intracellular replication (Strassmann and Shu, 2017). *Legionella* infections are a global public health concern presenting as either a severe pneumonia known as Legionnaires’ disease or Pontiac fever, a self-limiting flu-like syndrome (Herwaldt and Marra, 2018; Joseph *et al*., 2010; Lam *et al*., 2011; Wolter *et al*., 2016). Importantly, recent surveillance studies have indicated a steady increase of legionellosis incidences globally (Beauté and Network, 2017; Parr *et al*., 2015).

*L. pneumophila* infection in humans is considered to be the result of accidental environmental exposure and the selection for pathogenic traits among *L. pneumophila* is likely to be driven by co-selective pressures that exist in its natural habitat (O’Connor *et al*., 2011). The pivotal mechanism required for intracellular replication is the type IV secretion system (T4SS) that is conserved across all known members of the genus *Legionella* (Burstein *et al*., 2016). A very large repertoire of effector proteins in different combinations are encoded by *Legionella* species and can be secreted by this system to mediate critical host-pathogen interactions (Gomez-Valero *et al*., 2019). In addition, the ability to infect eukaryotic cells has evolved independently many times (Gomez-Valero *et al*., 2019). Despite this, less than half of all currently described *Legionella* species have been reported to cause human disease (Newton *et al*., 2010). Furthermore, there is an over-representation of a single serogroup (Sg-1) of *L. pneumophila* in human infections, which is responsible for more than 85% of all reported cases of legionellosis (Yu *et al*., 2002). Sg-1 strains can be further subdivided phenotypically using monoclonal antibodies (mAbs) that recognise various components of the lipopolysaccharide (LPS). The most prevalent mAb subtype in human infections is associated with an LPS *O*-acetyltransferase enzyme encoded by the *lag-1* gene (*lpg0777*), which confers an LPS epitope recognised by the mAb 3/1 from the Dresden mAb panel (Ditommaso *et al*., 2014; Kozak-Muiznieks *et al*., 2014).

Recently, it was shown that a very limited number (n=5) of *L. pneumophila* sequence types (ST)s are responsible for almost half of all human infections and that these clones have undergone very recent emergence and expansion (David *et al*., 2016). In addition, it was shown that recombination between the dominant STs has led to sharing of alleles that may be beneficial for human pathogenicity (David *et al*., 2016). However, the genetic basis for the enhanced human pathogenic potential of these STs is unknown. Early studies identified the capacity for some *L. pneumophila* strains to resist killing in human serum, a phenotype that may correlate with increased virulence (Plouffe *et al*., 1985). However, the molecular mechanism of strain-dependent serum resistance by *L. pneumophila* has remained elusive for over 30 years.

Here, we have carried out a population genomic analysis of a diverse dataset of 900 whole genome sequences of *L. pneumophila* isolates from both clinical and environmental sources to investigate the diversity of genotypes associated with human disease. Further, we employ genome-wide association analyses to identify genetic traits associated with human pathogenic strains revealing *lag-1* to be the most strongly associated determinant of human pathogenic potential in *L. pneumophila*. We demonstrate that acquisition of *lag-1* has occurred widely across the species by recent horizontal gene transfer and recombination, leading to strains that have enhanced resistance to serum killing and neutrophil phagocytosis. Importantly, functional analysis reveals that *lag-1* confers the capacity to evade human complement-mediated killing, suggesting a key role in the early stages of pathogenesis.

## Results and Discussion

### The potential for human infection is distributed across the *L. pneumophila* species phylogeny

In order to examine the diversity of *L. pneumophila* associated with human clinical infection in comparison to those from environmental sources, we carried out whole genome sequencing of 400 clinical and environmental *L. pneumophila* subsp. *pneumophila* isolates from an archived collection (1984 to 2015) of *Legionella* species held at the Scottish Haemophilus, Legionella, Meningococcus & Pneumococcus Reference Laboratory, Scottish Microbiology Reference Laboratory, Glasgow (SHLMPRL) as detailed in the SI Methods section. The dataset included 166 clinical isolates of infected patients, primarily from sputum and bronchoalveolar lavages (BAL) and 239 environmental isolates of *L. pneumophila* subsp. *pneumophila* from water sources such as cooling towers and plumbing systems sent to the reference laboratory for routine testing and surveillance. To place the SHLMPRL isolates into context of the known global diversity of *L. pneumophila* subsp. *pneumophila* isolates, we included 500 assembled whole genomes that were available in the public database (NCBI Genbank). We constructed a Maximum-Likelihood phylogenetic tree from 139,142 core genome single nucleotide polymorphisms (SNP) which indicates the segregation of the *L. pneumophila* subsp. *pneumophila* population into seven major clades (Fig. 1) each supported by a minimum of one and a maximum of two sub-clusters defined by Bayesian Analysis of Population Structure (BAPS) analysis (Supplemental Fig. 1). Isolates from the SHLMPRL collection were distributed among all major clades indicating that the isolates are representative of the global species diversity (Fig. 1). Although recombination has played an important role in diversification of the species, complicating the accurate reconstruction of the phylogeny (David *et al*., 2017), the major phylogenetic clusters are consistent with those identified in previous population studies employing fewer isolates (Qin *et al*., 2016; Underwood *et al*., 2013). Recombination is more likely between phylogenetically related (and genetically similar) isolates which may amplify the phylogenetic signal that defines these major groups (Skippington and Ragan, 2012). Of the 166 clinical isolates, 80 belonged to the five most common sequence types (ST) implicated in human infections (ST1, 23, 36/37, 47 and 62). The remaining 86 clinical isolates were from STs distributed across all seven major phylogenetic clusters (Fig. 1). These data indicate that although only 5 STs are responsible for almost half of all infections, the other half of human infection potential comes from diverse genetic backgrounds distributed across the species. Of note, five clusters from distinct outbreaks of Legionnaires’ disease in Scotland between 1985 and 2012 originated from different clades (Fig. 1). Similarly, the 239 environmental isolates are also distributed across all major phylogenetic groups (Fig. 1). Notably, 34% (81 of 239) of environmental isolates belong to one of the five clinically dominant STs suggesting that at least a third of all environmental *L. pneumophila* in Scotland have human pathogenic potential. Overall, although some STs appear to have higher human pathogenicity, our findings support the understanding that the capacity for causing human disease is widely distributed across diverse *L. pneumophila* genetic backgrounds.

**Fig. 1.**
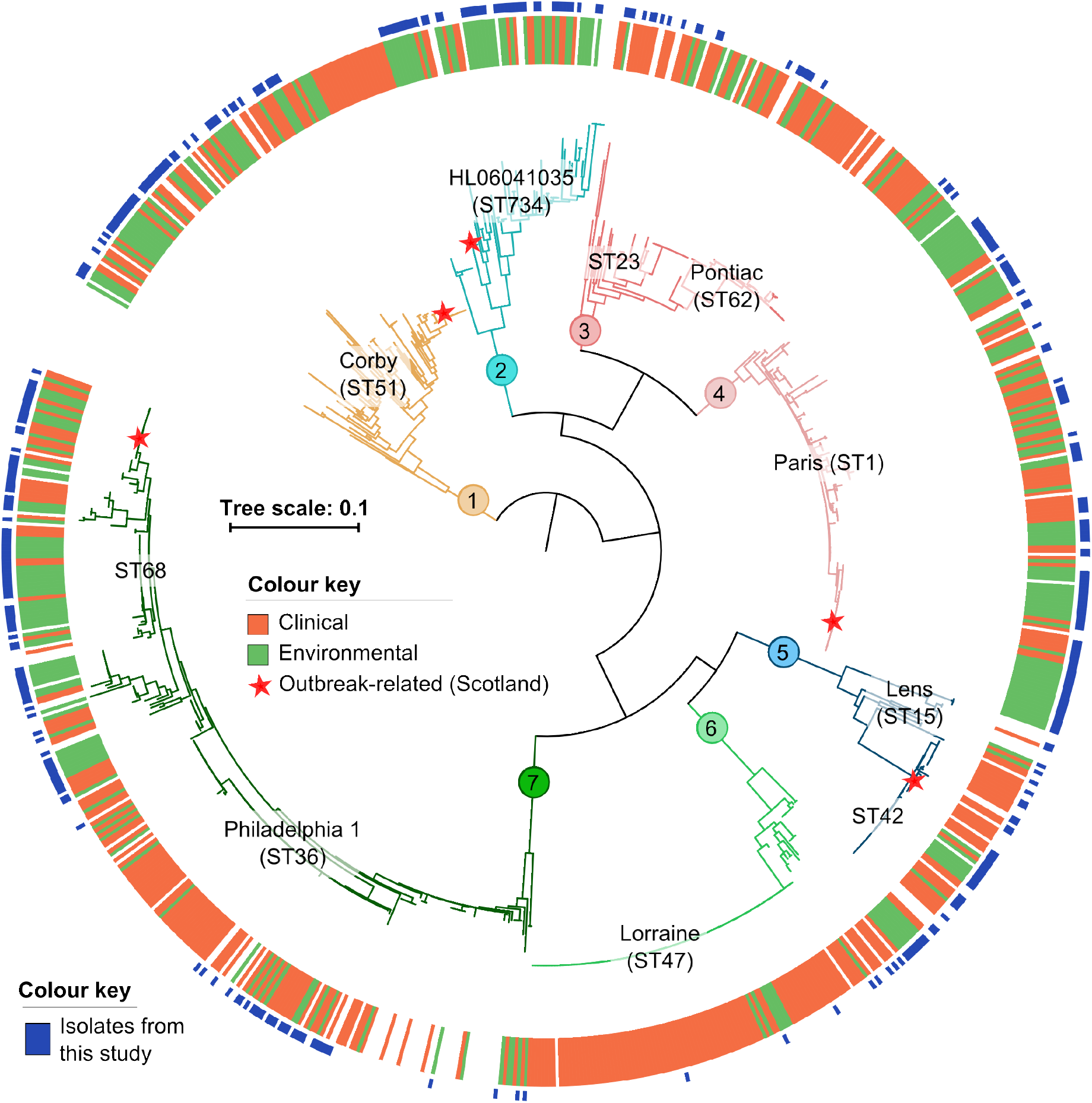
Human clinical isolates of *L. pneumophila* are widely distributed across the global species diversity. A maximum-likelihood phylogeny of 900 isolates based on 139,142 core genome SNPs divides the subspecies into seven major clades that are also supported by BAPS clustering. Isolates linked to five major outbreaks that have occurred in Scotland, UK between 1985 and 2012 originated from different lineages are indicated. In addition, isolates associated with sporadic cases of human infection have also emerged from diverse genetic backgrounds across all major clades. For reference, the position of five major disease associated clones (ST1, 23, 36/37, 47 and 62) and well-characterised reference genomes are indicated.

### Genome-wide association analysis of *L. pneumophila* reveals genetic traits associated with human pathogenicity

The factors that contribute to the enhanced human pathogenic capacity of some *L. pneumophila* clones are unknown. Our strategy, to address this gap in understanding, involved sequencing of large numbers of genetically diverse environmental isolates, affording for the first time, a large-scale genome-wide association study (GWAS) of clinical vs environmental isolates to explore the bacterial genetic basis for human clinical disease. Initially, we performed systematic subsampling on our dataset to reduce the number of closely related isolates originating from the same type of source, either clinical or environmental, while retaining genetic diversity. This step removes over-represented endemic or epidemic clonal lineages in the dataset that arise from opportunistic, convenience sampling and also represents an additional control for a stratified bacterial population structure. From this reduced dataset (n=452), we used the programme SEER to identify a total of 1737 *k-mers* that were enriched (significantly associated, p < 0.05) among clinical isolates in comparison to environmental isolates. Mapping to the *L. pneumophila* Philadelphia 1 reference genome (Accession number: AE017354) revealed that 39% (n=673) of the *k*-mers aligned to a region of the genome spanning between loci *lpg0748* and *lpg0781* representing an 18 kb cluster of genes involved in LPS biosynthesis and modification (Fig. 2) (Petzold *et al*., 2013). A total of 22 genes in this cluster each had at least one significantly enriched *k*-mer that mapped to it: namely, *lpg0751, lpg0752, lpg0755, lpg0758, lpg0759, lpg0760, lpg0761, lpg0762, lpg0766, lpg0767, lpg0768, lpg0769, lpg0771, lpg0772, lpg0773, lpg0774, lpg0775, lpg0777 (lag-1), lpg0778, lpg0779, lpg0780* and *lpg0781* (Fig. 2, Supplemental Table 2). Two additional conservative SEER analyses using more stringently subsampled datasets resulted in a smaller number of significantly enriched *k*-mers (205 and 61, respectively), that converge on a single gene within the LPS cluster (*lpg*0777, *lag-1*) that encodes an *O*-acetyltransferase involved in modification of the O-antigen of the *L. pneumophila* Sg-1 LPS (Supplemental Fig. S2). To corroborate the initial findings of the *k*-mer based SEER approach, we employed a different GWAS method (SCOARY) that examined the distribution of orthologous genes using the pan-genome pipeline ROARY (Brynildsrud *et al*., 2016; Page *et al*., 2015). Consistent with SEER, this approach also indicated that the LPS biosynthesis genes were enriched among clinical isolates with *lag-1* exhibiting the strongest statistical support over other Sg-1-associated LPS genes (Supplemental Table S2). The corrected (Benjamini-Hochberg) p-value for *lag-1* is several orders of magnitude lower (9.74E-11) than other Sg-1 LPS genes that were above the significance threshold (*lpg0779:* 2.35E-05, *lpg0780:* 2.35E-05, *lpl0815/lpg0774:* 3.84E-05 and *lpg0767*: 6.92E-05). As mentioned, *lag-1* has previously been reported to be prevalent among clinical isolates (Edelstein and Edelstein, 1993; Lück *et al*., 2001; Lück *et al*., 2002; Lüneberg *et al*., 1998; N. Whitfield and S. Swanson, 2006). However, our large pangenome-wide analysis of the *L. pneumophila* species indicates that of all 11198 accessory genes, *lag-1* has the strongest association with clinical isolates, suggesting a pivotal role in human disease.

**Fig. 2.**
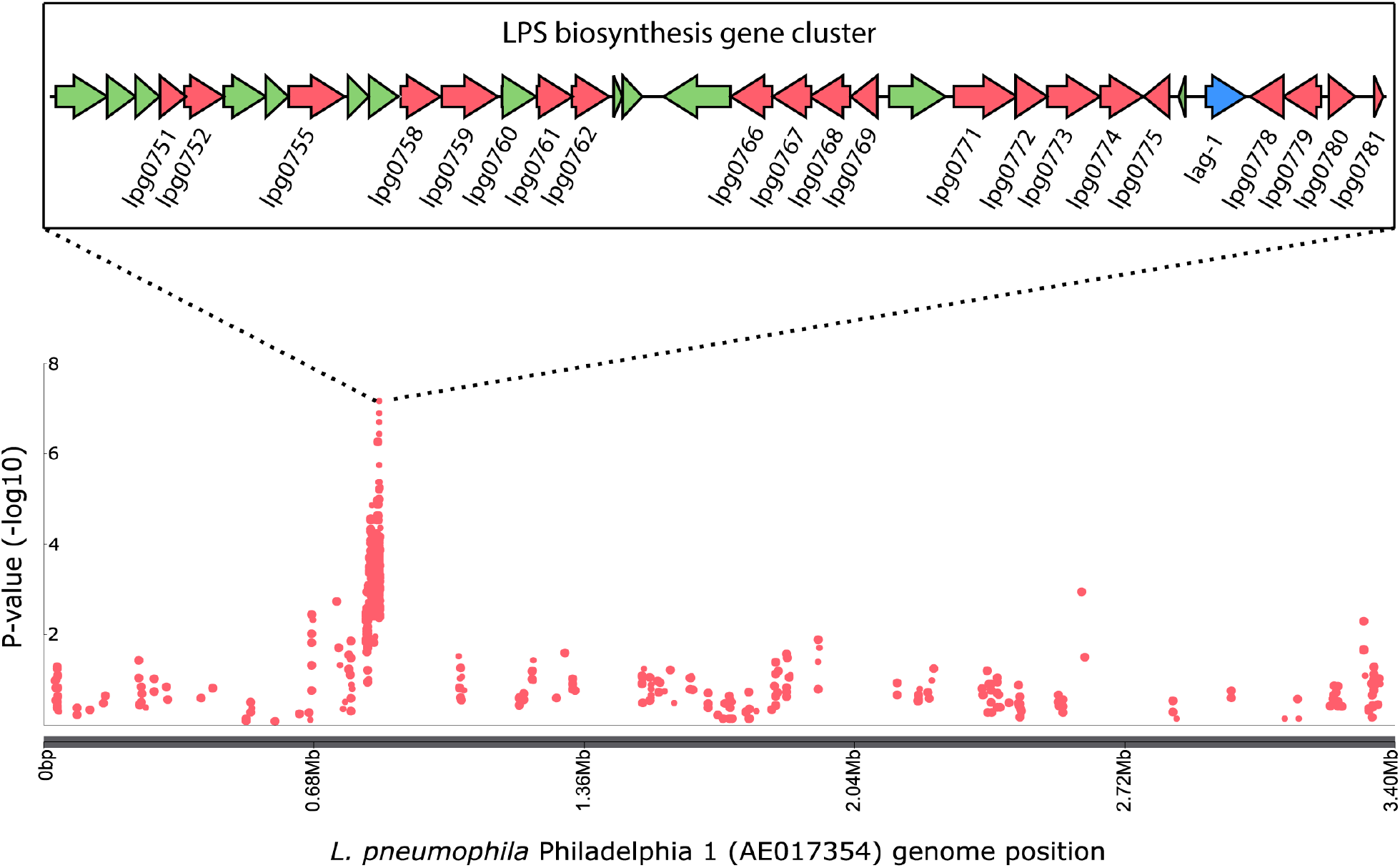
GWAS reveals gene sequences associated with clinical isolates of *L. pneumophila*. Manhattan plot showing the genomic position of *k*-mers significantly associated with clinical isolates. The LPS biosynthesis and modification locus (822,150bp–855,010bp: *lpg0751-lpg0781*) in the Philadelphia 1 reference genome (AE017354) is represented and genes with significant *k*-mer associations coloured in red, with *lag-1* in blue.

Sg-1 LPS genes were found in high frequency throughout the population, and in each major lineage indicating species-wide gene transfer. The Sg-1 LPS cluster is also found in other subspecies of *L. pneumophila* that can infect humans, such as subspecies *fraseri* and *pascullei* but has not been reported in other species of *Legionella* (George *et al*., 2016; Kozak-Muiznieks *et al*., 2016) Here, we observed that *lag-1* can be associated with different combinations of Sg-1 LPS genes and still express the expected *lag-1* phenotype represented by the mAb 3/1-specific epitope (Dresden mAb scheme) (Supplemental Table S1) (Helbig *et al*., 2002; Helbig *et al*., 1995). However, genetic instability and phase variation has been reported to affect the mAb phenotype between closely-related strains (Amemura-Maekawa *et al*., 2012; Bernander *et al*., 2003). One mechanism of phase variation is the excision of a 30 kb genetic element that results in a change in mAb specificity manifested by a loss of virulence in guinea pigs and loss of resistance to complement (Kooistra *et al*., 2002; Luneberg *et al*., 2001).

### Recombination has mediated the dissemination of three dominant *lag-1* alleles across the *L. pneumophila* species

It has been reported since the 1980s that isolates expressing the epitope recognized by the mAb 3/1 are more frequently associated with isolates from community-acquired and travel-associated infections (Harrison *et al*., 2007; Helbig *et al*., 2002; Kozak *et al*., 2009). However, the pathogenic basis for this association remains a mystery. In order to further investigate the role of the clinically-associated gene *lag-1*, we examined its diversity and distribution across the 900 *L. pneumophila* genomes employed in the current study. In total, three major allelic variants of *lag-1* that had been previously identified to be representative of reference strains, Philadelphia, Arizona, and Corby, respectively, were identified (Kozak *et al*., 2009). In our data set, variant 1 (Philadelphia), was present in 195 (22%), variant 2 (Arizona) in 195 (22%) and variant 3 (Corby) in 178 (20%) of the 900 isolates examined (Fig. 3A). Each variant was distributed across the phylogeny, with variant 1 found in 3 major clades (1, 4 and 7) and variants 2 and 3 identified among isolates of all 7 major clades (Fig. 3A). Of note, clade 2 has the lowest frequency (12%) of isolates encoding the *lag-1* gene and is also characterized by an under-representation of clinical isolates (31%). In addition to the three major *lag-1* alleles, we identified a relatively small number (n=40) of derived minor allelic variants that differ by <1% nucleotide identity from any of the 3 major variants (Supplemental Table S3). Of these, only 10 are predicted to encode for full-length proteins suggesting most are likely to be non-functional pseudogenes. The limited number of allelic variants of *lag-1* and their broad distribution across the species phylogeny indicates frequent horizontal dissemination of recently acquired *lag-1* alleles driven by a strong selection pressure for *lag-1* function. The closest homolog of *lag-1* in the NCBI non-redundant protein database shares only 45% amino acid sequence identity and encodes a putative acyltransferase present in an environmental species of the genus *Pseudomonas*. The selective advantage for *L. pneumophila* to maintain *lag-1* in the environment is still unclear. It has been proposed that the increased hydrophobicity of LPS when acetylated by *O*-acetyltransferase may enhance *L. pneumophila* survival in amoebae vacuolar compartments (Fernandez-Moreira *et al*., 2006).

**Fig. 3.**
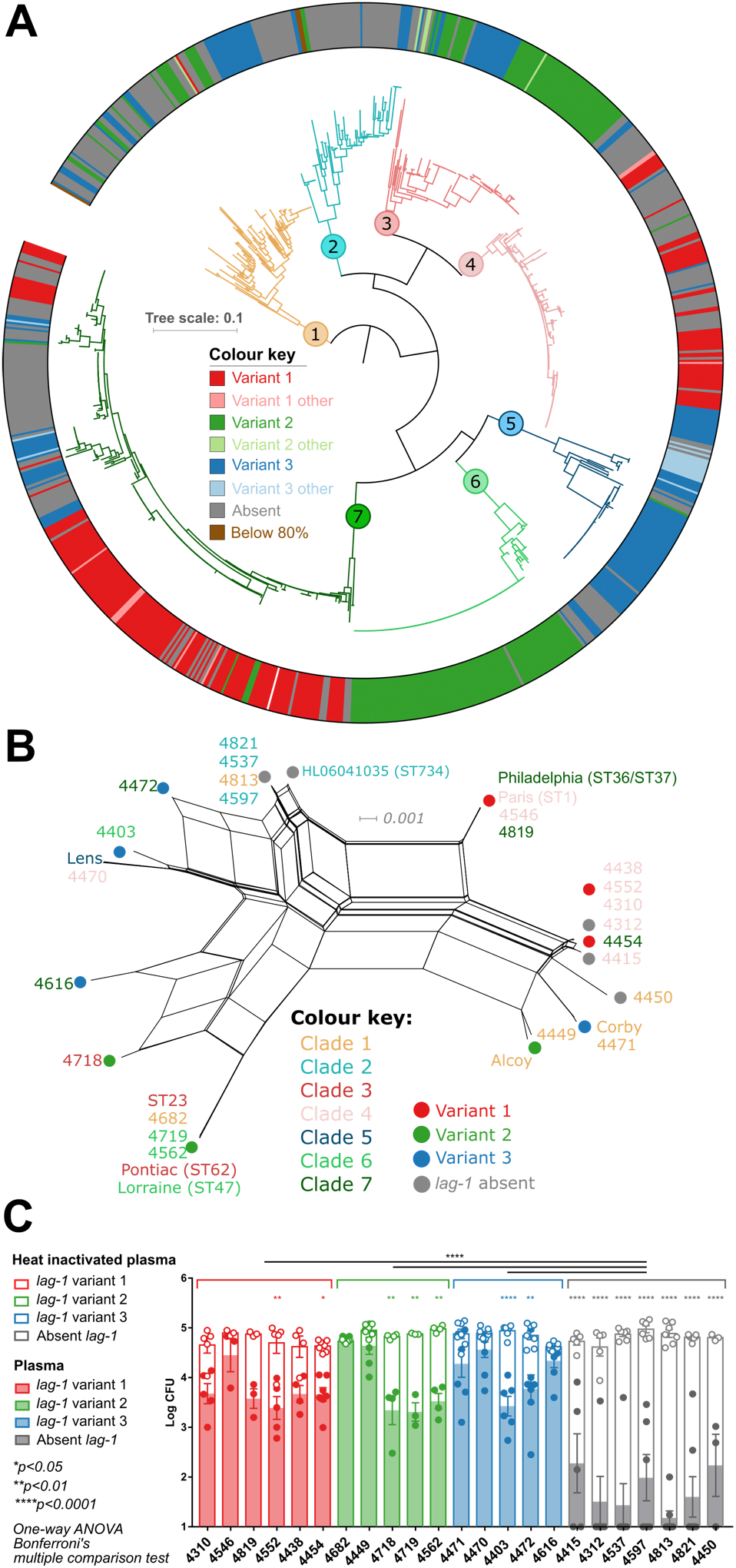
*lag-1* variants have disseminated horizontally across the species correlating with resistance to human plasma killing. **A) Core-genome SNP-based phylogenetic tree indicating the distribution of the 3 major allelic variants of *lag-1* across Sg-1 strains.** Variant 1 (Philadelphia, Red), Variant 2 (Arizona, Green) and Variant 3 (Corby, Blue). Each major clade is associated with at least two different *lag-1* alleles. Minor alleles that are within >99% nucleotide sequence identity to a major variant are shown with lighter colours. Isolates not encoding *lag-1* are coloured grey. **B) A neighbour-net phylogenetic network indicating recombination of Sg-1 genes including *lag-1*.** The network was drawn using uncorrected P-distances with the equal angle method in Splitstree from a concatenated alignment of 10 conserved Sg-1 genes that were significantly associated with clinical isolates (9405 positions). Orthologs of 10 LPS cluster genes were extracted from 16 phylogenetically representative isolates coding 3 different variants of *lag-1*, 7 isolates missing *lag-1* and 9 reference genomes (Lens, HL06041035, Philadelphia-1 [ST36], Paris [ST1], Corby, Alcoy, Pontiac [ST62], Lorraine [ST47], ST23). **C) Presence of *lag-1* correlates with resistance to killing in human plasma**. *L. pneumophila* isolates containing the allelic Variant 1, 2, and 3 of *lag-1* (depicted in red, green, and blue, respectively) or do not encode *lag-1* (depicted in grey) were incubated with human plasma (coloured bars and dots) or heat inactivated plasma (open bars and dots) for 1 hour at 37°C. Each coloured or open dot per bar represents the average count of an individual plasma donor.

Horizontal transfer of the 18 kb LPS biosynthesis gene cluster between Philadelphia-1 (ST36) and Paris (ST1) has been reported previously (Cazalet *et al*., 2008). To investigate the potential role of recombination in the distribution of the LPS genes including *lag-1*, we carried out a split network analysis based on the concatenated alignment of 10 LPS biosynthesis genes that were present in 23 representative isolates from across the phylogeny encoding different *lag-1* variants (Fig. 3B). This analysis revealed extensive reticulation consistent with recombination across the gene cluster and identified horizontal transfer of LPS genes between three major clinical sequence types, ST23, ST47 and ST62, respectively (Fig. 3B). This network analysis revealed that even highly similar LPS biosynthesis gene clusters, such as those found in isolates 4454, Corby and Alcoy or 4616 and 4718, encode different variants of *lag-1*, consistent with recent gene conversion of *lag-1*. We also identified three examples of closely related genomes (average nucleotide identity across the genome of > 99.8%) that exhibited a signature of homologous recombination affecting *lag-1* and the surrounding genomic region (Supplemental Fig. S3). These findings expand on a previously proposed mechanism of *lag-1* gene deletion by Kozak and colleagues (Kozak *et al*., 2009) and demonstrate that recombination can restore a disrupted or missing *lag-1* gene (Supplemental Fig. S3A and B) or replace one functional *lag-1* variant with another (Supplemental Fig. S3C).

A comparison of re-assortment rates across major phylogenetic lineages showed no significant differences between LPS and non-LPS genes across the *L. pneumophila* genome (*p*=0.6275), suggesting that recombination is active across the genome (Supplemental Fig. S4 and Supplemental Table S4). Taken together, these findings highlight the wide dissemination of *lag-1* and other LPS genes by recombination.

### *lag-1* confers resistance to killing by human plasma and broncho-alveolar lavage

Previous studies have reported an epidemiological correlation between *lag-1* or its mAb 3/1-recognised epitope and clinical disease (Edelstein and Edelstein, 1993; Lück *et al*., 2001; Lück *et al*., 2002; Lüneberg *et al*., 1998; N. Whitfield and S. Swanson, 2006) but an understanding of a role in pathogenesis has proved elusive. Of note, an early study described an increased ability of an endemic nosocomial *L. pneumophila* strain to survive complement killing when compared to an environmental strain collected from the same medical facility (Plouffe *et al*., 1985). To investigate this unexplained phenotype in the context of *lag-1* genotype, we examined resistance to killing in human plasma for the aforementioned 23 *L. pneumophila* strains selected to represent the diversity of *lag-1* genotypes from across the phylogeny. We observed that all strains lacking a *lag-1* gene were susceptible to killing in plasma, whereas the majority (9 of 16) of isolates containing *lag-1* exhibited enhanced resistance, independent of the *lag-1* variant encoded (Fig. 3C).

Within our dataset, we identified a group of epidemiologically-related isolates from a single Scottish healthcare facility typed as ST5 (a single locus variant of ST1) which were predicted to vary with regard to *lag-1* functionality. ST5 has ony been found in this location, to date. The earliest isolates obtained from a nosocomial outbreak in 1984/1985 (Macfarlane and Worboys, 2012) contained variant 3 (Corby) of the *lag-1* gene and all were found to express the mAb 3/1 epitope (Supplemental Table S1). In contrast, 16 of 19 environmental isolates from the same healthcare facility 12 to 21 years later (1997 to 2006) contained multiple independent mutations in *lag-1* predicted to disrupt functionality. Specifically, a nonsense mutation (L48*), an insertion of a transposase, and the acquisition of a deleterious substitution (H28L) (Fig. 4A), each correlated with a lack of reactivity with mAb 3/1 from the Dresden panel classification (Supplemental Table S1). Consistent with our previous findings (Fig. 3C), the presence of a functional LPS *O*-acetyltransferase in this epidemiological cluster also correlated with resistance to killing in human plasma, whereas isolates with a non-functional *lag-1* were susceptible to killing, independent of the type of deactivating mutation (Fig. 4B). Of note, no clinical episodes of disease were identified to be caused by this epidemiological cluster after 1985, and all clinical isolates contained a functional *lag-1* gene. The identification of multiple independent mutations associated with loss of *lag-1* function suggests a selection pressure that drives the inactivation of the *lag-1* gene and LPS *O*-acetylation in this environment. The trend of *lag-1* gene loss or deactivation was not observed among longitudinal ST1 healthcare facility-associated isolates from a cluster sequenced in a recent study by David and colleagues (David *et al*., 2017). However, a study on starvation of *L. pneumophila* in ultrapure water showed that in a short-term period, the viable cell numbers of all mAb 3/1-positive strains decreased strongly compared to the other strains suggesting a negative selection for *lag-1* function in some water environments (Schrammel *et al*., 2018). Overall, these data indicate that the presence of a functional *lag-1* correlates with enhanced resistance to serum killing.

**Fig. 4.**
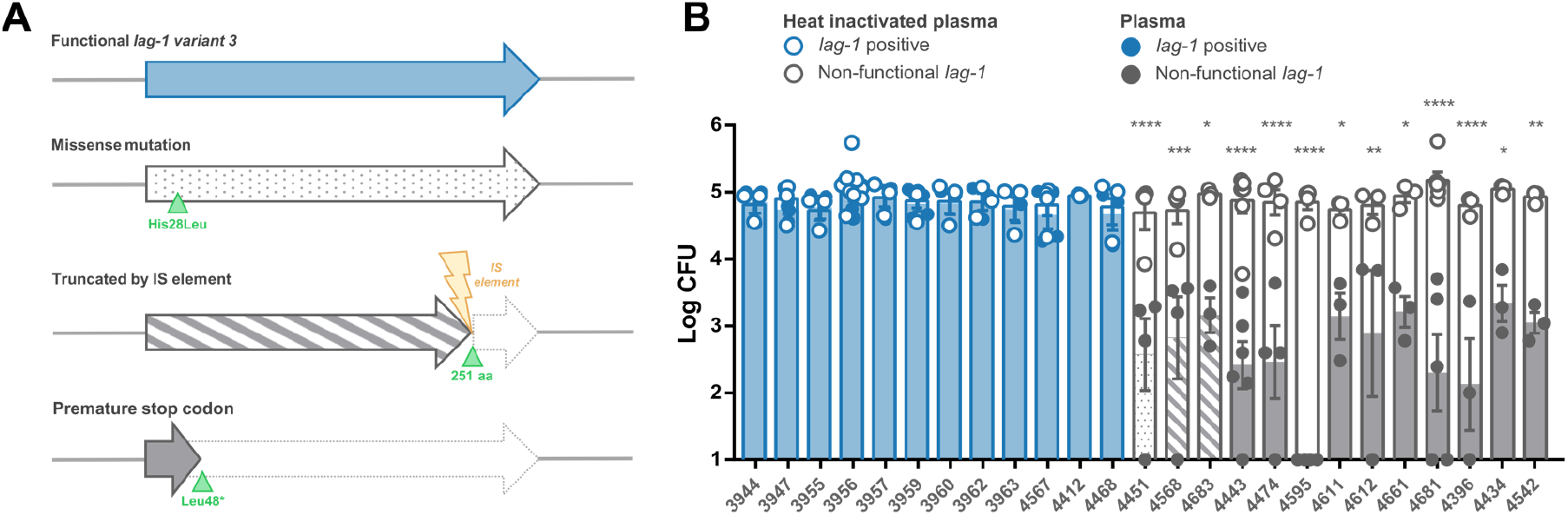
Multiple natural independent deactivating mutations in *lag-1* gene conferred susceptibility to complement killing. **A) Schematic representation of the natural *lag-1* deactivating mutations.** In blue - functional *lag-1*, with dots - *lag-1* with His28Leu mutation, with lines - *lag-1* truncated by an insertion sequence in aa 251, in grey - *lag-1* truncated by a premature stop codon. **B) Isolates with *lag-1* mutations show increased susceptibility to killing in human plasma.** Closely related isolates from the same hospital-associated cluster were incubated with human plasma (coloured and patterned bars and dots) or heat inactivated plasma (open bars and dots) for 1 hour at 37°C. Each dot represent an average count of an individual plasma donor. One-way ANOVA, Tukey’s multiple comparisons test.

To investigate the role of *lag-1* in mediating resistance to serum killing, we introduced the *lag-1* gene encoded on an expression plasmid (pMMB207) into two *L. pneumophila* mAb 3/1 epitope-negative strains. We introduced the three *lag-1* variants into the strain 4681 that contains a non-functional *lag-1* gene due to the insertion of a transposase, and the *lag-1* variant 1 into the *lag-1* negative strain 4312 which contains a 1 bp deletion resulting in a frameshift at position 156. Introduction of the plasmid resulted in *lag-1* expression and LPS 8-*O*-acetylation as confirmed by flow cytometric detection of the mAb 3/1 epitope (Fig. 5A). Strikingly, complementation of the strains with *lag-1* resulted in resistance to killing in human serum independently of the *lag-1* variant (Fig. 5B). Associated with the resistance to serum killing, we observed a decrease in human C3 deposition at the surface of the *lag-1* expressing bacteria when compared to the isogenic *lag-1* negative strain, at levels similar to the wild-type *lag-1* positive isolate 3656 (Supplemental Fig. S5). Of note, it was previously reported that a wild-type *lag-1* positive isolate and a spontaneously derived *lag-1* mutant demonstrated equivocal levels of resistance to serum killing (Lück *et al*., 2001). We speculate that the discordance with our findings could be due to phase-variable expression of LPS (Lüneberg *et al*., 1998) or differences in transcriptional levels of *lag-1* gene that have been previously observed depending on culture conditions (Faucher *et al*., 2011).

**Fig. 5.**
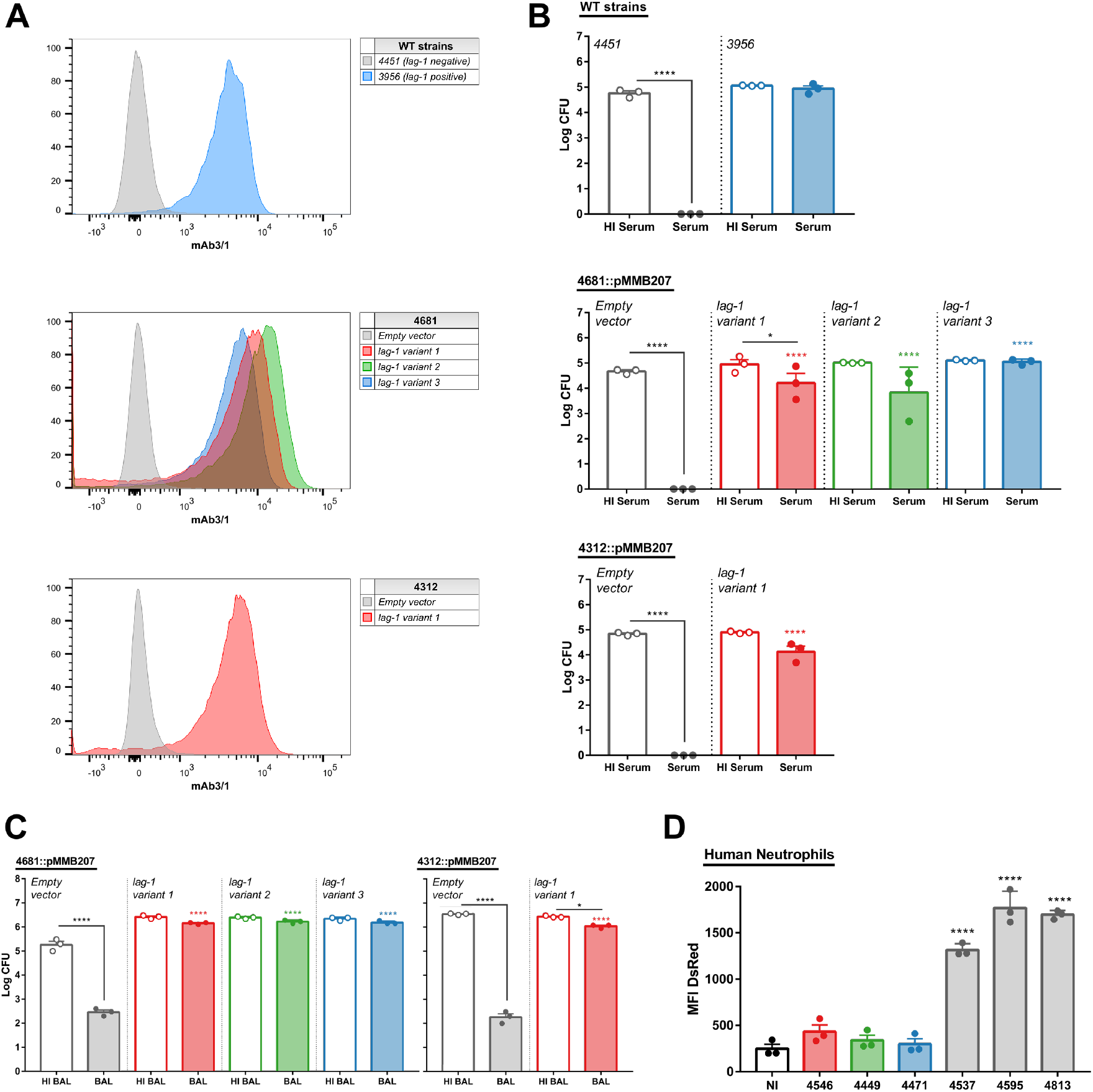
*lag-1* confers resistance to complement-killing and neutrophil internalization of *L. pneumophila*. **A) Introduction of any *lag-1* gene variant leads to mAB 3/1 epitope expression.** Detection of mAb 3/1 epitope by flow cytometry in WT and isogenic mutants of *L. pneumophila* isolates expressing the *lag-1* variant 1, 2 or 3. **B) *lag-1* expression confers serum complement resistance to *L. pneumophila* strains with non-functional *lag-1* gene.** Isolates were incubated with human serum or heat inactivated serum for 1 hour at 37°C. Each point represents an average of triplicate CFU counts of a single sera donor. Bars represent mean+SEM. One-way ANOVA, Tukey’s multiple comparisons test, *p<0.05, ****p<0.0001. Comparasion to *Empty plasmid* isogenic strain represented on the top of the bars. **C) mAb 3/1 negative *L. pneumophila* strains are susceptible to bovine BAL complement killing.** Isolates were incubated with concentrated bovine BAL or heat inactivated BAL for 1 hour at 37°C. Each point represents a replicate. Bars represent mean+SEM. One-way ANOVA, Tukey’s multiple comparisons test, *p<0.05, ****p<0.0001. Comparasion to *Empty plasmid* isogenic strain represented on the top of the bars. **D) *lag-1* confers resistance to human neutrophil phagocytosis.** WT *lag-1* positive and negative strains expressing DsRed fluorescent protein were pre-incubated with 10% human serum 15 min prior incubation with human neutrophils for 30 min. Phagocytosis was evaluated by measuring neutrophils DsRed fluoresce by flow cytometry. Data representative of two independent experiments. *L. pneumophila* strains expressing *lag-1* variant 1, 2 or 3 are represented in red, green or blue, respectively. Strains that do not express a functional *lag-1* gene are represented in gray. Negative controls are represented in black. Bars represent mean+SEM. One-way ANOVA, Dunnett’s multiple comparisons test to non-infected control, *p<0.05, ****p<0.0001. HI - heat inactivated, BAL – bronchoalveolar lavage, NI – non-infected.

Classical pathway complement activity exists in healthy human broncho-alveolar lavage (BAL), despite the relatively low concentration of some complement proteins (Watford *et al*., 2000), and exposure to aerosolized LPS leads to a rapid increase of the level of these proteins in the lung of human volunteers (Bolger *et al*., 2007). The importance of this innate immune mechanism in lung health is supported by the observation that many patients with deficiencies in complement proteins or complement receptors have recurrent respiratory infections (Figueroa and Densen, 1991). Here, we examined the role of *lag-1* in resistance to killing in bovine BAL. Similar to the impact of human serum, incubation of *lag-1* negative *L. pneumophila* with concentrated bovine BAL resulted in significant bacterial killing compared with heat inactivated fluid (Fig. 5C), and complementation with each *lag-1* variant conferred resistance to killing (Fig. 5C). Taken together, our data reveal a key role for *lag-1* in conferring resistance to complement killing in both serum and bronchoalveolar fluid.

### Presence of lag-1 correlates with resistance to neutrophil phagocytosis

The role of the complement system in the lung immune defences extends beyond the proteolytic cascade associated with bacterial lysis, as opsonization with complement C3b protein induces phagocytosis of the opsonized targets by neutrophils and macrophages (Heesterbeek *et al*., 2018). Accordingly, to test the effect of *lag-1* expression on phagocytosis of *L. pneumophila* by neutrophils, human blood purified neutrophils were incubated with *L. pneumophila* fluorescent strains in the presence of non-immune human serum (Fig. 5D).

We selected *L. pneumophila* strains from different phylogenetic groups that express the three *lag-1* variants, in addition to three *lag-1* negative strains. We observed a considerably higher mean fluorescence intensity (MFI) in neutrophils infected with *lag-1* negative strains compared to lag-1 positive strains, indicating reduced adherence or phagocytosis that correlated with lag-1 expression (Fig. 5 D; Fig. S6). Therefore, the ability of *lag-1* positive *L. pneumophila* strains to escape complement deposition correlates with the capacity to escape internalization and killing by human phagocytes. These data are consistent with early *L. pneumophila* studies, where *L. pneumophila* strain Philadelphia 1 (*lag-1* positive) was demonstrated to be resistant to complement and neutrophil killing in the absence of specific antibodies (Horwitz and Silverstein, 1981; Verbrugh *et al*., 1985). There is increasing evidence for the pivotal role of neutrophils in the resolution of *L*. *pneumophila* lung and either neutrophil depletion or blockage of recruitment to infected lungs renders mice susceptible to *L. pneumophila* (LeibundGut-Landmann *et al*., 2011; Mascarenhas *et al*., 2015; Tateda *et al*., 2001a; Tateda *et al*., 2001b; Ziltener *et al*., 2016). Taken together, our data indicate that *L. pneumophila lag-1* mediated inhibition of complement deposition, disrupts the innate immune response via inhibition of complement-mediated lysis and blocking recognition by phagocytes and subsequent internalization.

### Lag-1 confers resistance to killing by the classical complement pathway

We next investigated if a specific complement pathway was responsible for the serum killing of the Sg-1 mAb 3/1-negative *L. pneumophila*. As a similar inhibitory effect of *lag-1* on bacterial killing was observed in both serum and plasma, we employed serum for these experiments to facilitate the use of commercially available depleted serum samples. EDTA can be used to inhibit the classical, lectin and alternative pathways via chelation of both Ca^2+^ and Mg^2+^, whereas chelation with EGTA/Mg^2+^ inhibits the Ca^2+^-dependent classical and lectin pathways only, leaving the alternative pathway unaffected (Fig. 6A). Killing assays in human serum in the presence of either EDTA or EGTA/Mg^2+^ resulted in complete abrogation of *lag-1*-negative strains susceptibility to complement, and no difference in cell viability compared to incubation in heat inactivated serum, suggesting the alternative complement pathway is insufficient to kill *L. pneumophila* (Fig. 6B). To distinguish the role of the classical from the lectin pathway, killing assays were performed in the presence of mannose that competes for the association of mannose binding lectin (MBL) with the bacterial surface and blocks the lectin complement pathway (Moller-Kristensen *et al*., 2006). Using this approach, there was no effect on bacteria viability, indicating that the lectin pathway is not required for *L. pneumophila* Sg-1 killing, consistent with the previous report that MBL polymorphisms are not associated with a higher risk for legionellosis (Herpers *et al*., 2009). The exclusion of the lectin pathway suggests the essential role of the classical pathway in the complement killing of *L. pneumophila*. To confirm this finding we depleted serum of C1q, required for the classical pathway, resulting in loss of serum-mediated killing (Fig. 6B). It is noteworthy that we also observed an increase in bacterial viability in the presence of Factor B-depleted serum (Fig. 6B), implying a possible role for the alternative pathway in amplification of classical pathway activation as previous described for *Streptococcus pneumoniae* (Brown *et al*., 2002). Previously, purified *L. pneumophila* LPS was reported to activate both classical and alternate pathways, primarily through the activation of the classical pathway dependent on natural IgM antibodies (Mintz *et al*., 1992). In another study, complement C1q protein was demonstrated to bind to the major outer membrane protein (MOMP) of *L. pneumophila*, activating complement in an antibody-independent way (Mintz *et al*., 1995).

**Fig. 6.**
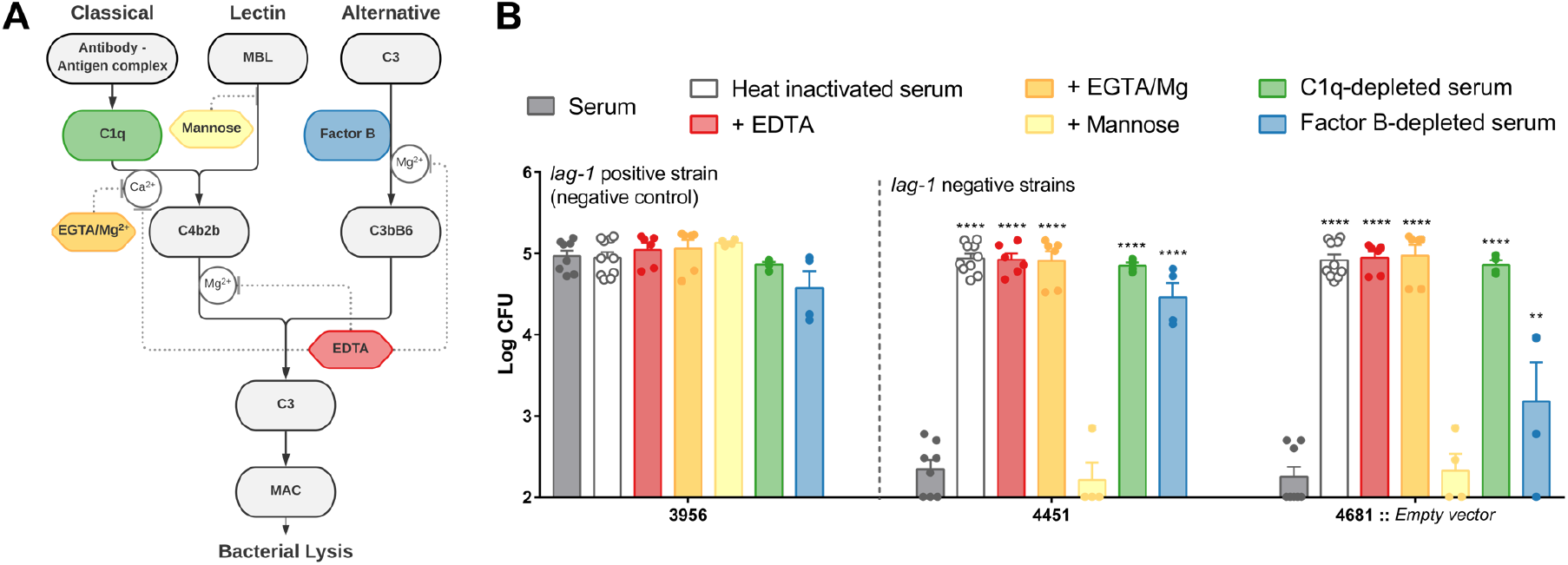
The classical pathway is essential for complement-mediated killing of *L. pneumophila*. **A) Simplified schematic representation of the complement pathways and the inibitors used in this study. B) Inhibiton of both classical and alternative pathways confers *L. pneumophila* resistance to complement killing in human serum.** 3956, 4451, 4681::*Empty plasmid* isolates were incubated with 90% heat inactivated serum, serum, serum with EDTA, EGTA/Mg^2+^ or mannose, Factor B- or C1q-depleted serum for a 1 hour at 37°C. Statistical analysis when compared to serum on top of the bars. One-way ANOVA, Tukey’s multiple comparisons test, **p<0.01 ****p < 0.0001

### Concluding comments

The steady global increase in *L. pneumophila* infections is worrisome and studies that explore the evolutionary basis of the increased pathogenicity of some clones can provide insights into the nature of the public health threat posed. Here we have employed a combined large-scale population level study of clinical and environmental isolates, and a functional *ex vivo* analysis to reveal the basis for serum resistance in *L. pneumophila*. The identification of a specific modification of LPS that is required for resistance to classical complement mediated killing and inhibits phagocytosis by neutrophils could inform the design of novel therapeutic approaches that subvert the capacity of *L. pneumophila* to resist innate immune killing during severe human infection.

## Materials and Methods

### Bacterial strains and plasmids

Bacterial strains and plasmids employed in this study are described in SI Appendix. *L. pneumophila* cells were cultured in yeast extract broth (YEB) with constant shaking at 37 °C in the presence or absence of chloramphenicol (SI Appendix, SI Materials and Methods).

### Gene cloning and complementation

The three *lag-1* gene variants were cloned into the vector pMMB207 and transformed by electrophoresis into the *L. pneumophila* isolate 4681 or 4312. pSW001 plasmid, contitutevely expressing DsRed was transformed into *L. pneumophila* as described in SI Materials and Methods (Chen *et al*., 2006). Expression of the mAb 3/1 epitope and DsRed fluorescence was confirmed by flow cytometry (SI Appendix, SI Materials and Methods).

### Whole genome sequence analysis

Sequence data are available on the European Nucleotide Archive, accession number PRJEB31628. For detailed methods please refer to SI Appendix, SI Materials and Methods.

### Complement killing assays

*L. pneumophila* was incubated for 1 hour at 37°C in 90% normal human serum/plasma, heat inactivated serum/plasma, bovine concentrated BAL or heat inactivated BAL. Inhibition of complement pathways was carried out by adding 12.5 mM EDTA (Sigma Aldrich), 12.5 mM EGTA/Mg^2+^ (Sigma Aldrich) or 100 mM mannose (Acros Organics) to normal human serum or by using C1q- and Factor B-depleted serum (Pathway Diagnostics). (SI Appendix, SI Materials and Methods).

### Neutrophil assay

*L. pneumophila* transformed with pSW001 plasmid was incubated with 10% normal human serum for 15 min prior to incubation with purified human blood neutrophils for 30 min at 37°C, 600 rpm. Bacterial internalization was evaluated by flow cytometry (SI Appendix, SI Materials and Methods).

## Acknowledgements

This work was supported by funding to JRF from the Chief Scientists Office Scotland (Grant No. ETM/421) and the Wellcome Trust Collaborative award (Grant No. 201531/Z/16/Z). Computing resources were supported in part by MRC CLIMB (Grant Number: MR/L015080/1). We are grateful to Carmen Buchrieser for providing plasmid pMMB207. Thanks also to Dr Kenneth Baillie and Dr Sara Clohisey for organising the human blood donation study and the volunteers from the Roslin Institute who provided blood samples for the serum and plasma killing assays.

## Notes

### Competing Interest Statement

The authors have declared no competing interest.

## References

Amemura-Maekawa, J., Kikukawa, K., Helbig, J.H., Kaneko, S., Suzuki-Hashimoto, A., Furuhata, K., Chang, B., Murai, M., Ichinose, M., Ohnishi, M., et al. (2012). Distribution of monoclonal antibody subgroups and sequence-based types among *Legionella pneumophila* serogroup 1 isolates derived from cooling tower water, bathwater, and soil in Japan. Appl Environ Microbiol. 78(12), 4263–4270. Published online 2012/04/06 DOI: 10.1128/AEM.06869-11.

Beauté, J., and Network, T.E.L.D.S. (2017). Legionnaires’ disease in Europe, 2011 to 2015. Euro Surveill. 22(27). DOI: 10.2807/1560-7917.ES.2017.22.27.30566.

Bernander, S., Jacobson, K., Helbig, J.H., Lück, P.C., and Lundholm, M. (2003). A hospital-associated outbreak of Legionnaires’ disease caused by *Legionella pneumophila* serogroup 1 is characterized by stable genetic fingerprinting but variable monoclonal antibody patterns. J Clin Microbiol. 41(6), 2503–2508.

Bolger, M.S., Ross, D.S., Jiang, H., Frank, M.M., Ghio, A.J., Schwartz, D.A., and Wright, J.R. (2007). Complement levels and activity in the normal and LPS-injured lung. Am J Physiol Lung Cell Mol Physiol. 292(3), L748–759. Published online 2006/10/27 DOI: 10.1152/ajplung.00127.2006.

Brown, J.S., Hussell, T., Gilliland, S.M., Holden, D.W., Paton, J.C., Ehrenstein, M.R., Walport, M.J., and Botto, M. (2002). The classical pathway is the dominant complement pathway required for innate immunity to *Streptococcus pneumoniae* infection in mice. Proc Natl Acad Sci USA. 99(26), 16969–16974. Published online 2002/12/11 DOI: 10.1073/pnas.012669199.

Brynildsrud, O., Bohlin, J., Scheffer, L., and Eldholm, V. (2016). Rapid scoring of genes in microbial pan-genome-wide association studies with Scoary. Genome Biol. 17(1), 238. Published online 2016/11/25 DOI: 10.1186/s13059-016-1108-8.

Burstein, D., Amaro, F., Zusman, T., Lifshitz, Z., Cohen, O., Gilbert, J.A., Pupko, T., Shuman, H.A., and Segal, G. (2016). Genomic analysis of 38 Legionella species identifies large and diverse effector repertoires. Nat Genet. 48(2), 167–175. Published online 2016/01/11 DOI: 10.1038/ng.3481.

Cazalet, C., Jarraud, S., Ghavi-Helm, Y., Kunst, F., Glaser, P., Etienne, J., and Buchrieser, C. (2008). Multigenome analysis identifies a worldwide distributed epidemic *Legionella pneumophila* clone that emerged within a highly diverse species. Genome Res. 18(3), 431–441. Published online 2008/02/06 DOI: 10.1101/gr.7229808.

Chen, D.Q., Huang, S.S., and Lu, Y.J. (2006). Efficient transformation of *Legionella pneumophila* by high-voltage electroporation. Microbiol Res. 161(3), 246–251. DOI: 10.1016/j.micres.2005.09.001.

David, S., Afshar, B., Mentasti, M., Ginevra, C., Podglajen, I., Harris, S.R., Chalker, V.J., Jarraud, S., Harrison, T.G., and Parkhill, J. (2017). Seeding and Establishment of *Legionella pneumophila* in Hospitals: Implications for Genomic Investigations of Nosocomial Legionnaires’ Disease. Clin Infect Dis. 64(9), 1251–1259. DOI: 10.1093/cid/cix153.

David, S., Rusniok, C., Mentasti, M., Gomez-Valero, L., Harris, S.R., Lechat, P., Lees, J., Ginevra, C., Glaser, P., Ma, L., et al. (2016). Multiple major disease-associated clones of *Legionella pneumophila* have emerged recently and independently. Genome Res. 26(11), 1555–1564. DOI: 10.1101/gr.209536.116.

Ditommaso, S., Giacomuzzi, M., Rivera, S.R., Raso, R., Ferrero, P., and Zotti, C.M. (2014). Virulence of *Legionella pneumophila* strains isolated from hospital water system and healthcare-associated Legionnaires’ disease in Northern Italy between 2004 and 2009. BMC Infect Dis. 14, 483. Published online 2014/09/05 DOI: 10.1186/1471-2334-14-483.

Edelstein, P.H., and Edelstein, M.A. (1993). Intracellular growth of *Legionella pneumophila* serogroup 1 monoclonal antibody type 2 positive and negative bacteria. Epidemiol Infect. 111(3), 499–502. Published online 1993/12/01.

Faucher, S.P., Mueller, C.A., and Shuman, H.A. (2011). *Legionella pneumophila* transcriptome during intracellular multiplication in human macrophages. Front Microbiol. 2, 60. Published online 2011/04/04 DOI: 10.3389/fmicb.2011.00060.

Fernandez-Moreira, E., Helbig, J.H., and Swanson, M.S. (2006). Membrane vesicles shed by *Legionella pneumophila* inhibit fusion of phagosomes with lysosomes. Infect Immun. 74(6), 3285–3295. DOI: 10.1128/IAI.01382-05.

Figueroa, J.E., and Densen, P. (1991). Infectious diseases associated with complement deficiencies. Clin Microbiol Rev. 4(3), 359–395.

George, F., Shivaji, T., Pinto, C.S., Serra, L.A.O., Valente, J., Albuquerque, M.J., Vicêncio, P.C.O., San-Bento, A., Diegues, P., Nogueira, P.J., et al. (2016). A large outbreak of Legionnaires’ Disease in an industrial town in Portugal. Revista Portuguesa de Saúde Pública. 34(3), 199–208. Published online 16 November 2016 DOI: 10.1016/j.rpsp.2016.10.001.

Gomez-Valero, L., Rusniok, C., Carson, D., Mondino, S., Pérez-Cobas, A.E., Rolando, M., Pasricha, S., Reuter, S., Demirtas, J., Crumbach, J., et al. (2019). More than 18,000 effectors in the *Legionella* genus genome provide multiple, independent combinations for replication in human cells. Proc Natl Acad Sci U S A. 116(6), 2265–2273. Published online 2019/01/18 DOI: 10.1073/pnas.1808016116.

Harrison, T.G., Doshi, N., Fry, N.K., and Joseph, C.A. (2007). Comparison of clinical and environmental isolates of Legionella pneumophila obtained in the UK over 19 years. Clin Microbiol Infect. 13(1), 78–85. DOI: 10.1111/j.1469-0691.2006.01558.x.

Heesterbeek, D.A.C., Angelier, M.L., Harrison, R.A., and Rooijakkers, S.H.M. (2018). Complement and Bacterial Infections: From Molecular Mechanisms to Therapeutic Applications. J Innate Immun. 10(5-6), 455–464. Published online 2018/08/28 DOI: 10.1159/000491439.

Helbig, J.H., Bernander, S., Castellani Pastoris, M., Etienne, J., Gaia, V., Lauwers, S., Lindsay, D., Lück, P.C., Marques, T., Mentula, S., et al. (2002). Pan-European study on culture-proven Legionnaires’ disease: distribution of *Legionella pneumophila* serogroups and monoclonal subgroups. Eur J Clin Microbiol Infect Dis. 21(10), 710–716. Published online 2002/10/18 DOI: 10.1007/s10096-002-0820-3.

Helbig, J.H., Lück, P.C., Knirel, Y.A., Witzleb, W., and Zähringer, U. (1995). Molecular characterization of a virulence-associated epitope on the lipopolysaccharide of *Legionella pneumophila* serogroup 1. Epidemiol Infect. 115(1), 71–78.

Herpers, B.L., Yzerman, E.P., de Jong, B.A., Bruin, J.P., Lettinga, K.D., Kuipers, S., Den Boer, J.W., van Hannen, E.J., Rijkers, G.T., van Velzen-Blad, H., et al. (2009). Deficient mannose-binding lectin-mediated complement activation despite mannose-binding lectin-sufficient genotypes in an outbreak of *Legionella pneumophila pneumonia*. Hum Immunol. 70(2), 125–129. Published online 2008/12/13 DOI: 10.1016/j.humimm.2008.11.002.

Herwaldt, L.A., and Marra, A.R. (2018). Legionella: a reemerging pathogen. Curr Opin Infect Dis. 31(4), 325–333. DOI: 10.1097/QCO.0000000000000468.

Hilbi, H., Hoffmann, C., and Harrison, C.F. (2011). *Legionella spp.* outdoors: colonization, communication and persistence. Environ Microbiol Rep. 3(3), 286–296. Published online 2011/03/21 DOI: 10.1111/j.1758-2229.2011.00247.x.

Horwitz, M.A., and Silverstein, S.C. (1981). Interaction of the Legionnaires’ disease bacterium (*Legionella pneumophila*) with human phagocytes. I. *L. pneumophila* resists killing by polymorphonuclear leukocytes, antibody, and complement. J Exp Med. 153(2), 386–397. Published online 1981/02/01 DOI: 10.1084/jem.153.2.386.

Joseph, C.A., Ricketts, K.D., and Infections, C.o.b.o.t.E.W.G.f.L. (2010). Legionnaires’ disease in Europe 2007–2008. DOI: doi:10.2807/ese.15.08.19493-en.

Kooistra, O., Zähringer, U., Lüneberg, E., Frosch, M., and Knirel, Y.A. (2002). Phase variation in *Legionella pneumophila* serogroup 1, subgroup OLDA, strain RC1 influences lipid A structure. In: Legionella, (American Society of Microbiology), pp. 68–73.

Kozak-Muiznieks, N.A., Lucas, C.E., Brown, E., Pondo, T., Taylor, T.H., Frace, M., Miskowski, D., and Winchell, J.M. (2014). Prevalence of sequence types among clinical and environmental isolates of *Legionella pneumophila* serogroup 1 in the United States from 1982 to 2012. J Clin Microbiol. 52(1), 201–211. Published online 2013/11/06 DOI: 10.1128/JCM.01973-13.

Kozak-Muiznieks, N.A., Morrison, S.S., Sammons, S., Rowe, L.A., Sheth, M., Frace, M., Lucas, C.E., Loparev, V.N., Raphael, B.H., and Winchell, J.M. (2016). Three Genome Sequences of *Legionella pneumophila subsp. pascullei* Associated with Colonization of a Health Care Facility. Genome Announc. 4(3). Published online 2016/05/07 DOI: 10.1128/genomeA.00335-16.

Kozak, N.A., Benson, R.F., Brown, E., Alexander, N.T., Taylor, T.H., Shelton, B.G., and Fields, B.S. (2009). Distribution of *lag-1* alleles and sequence-based types among *Legionella pneumophila* serogroup 1 clinical and environmental isolates in the United States. J Clin Microbiol. 47(8), 2525–2535. Published online 2009/06/24 DOI: 10.1128/JCM.02410-08.

Lam, M.C., Ang, L.W., Tan, A.L., James, L., and Goh, K.T. (2011). Epidemiology and Control of Legionellosis, Singapore. Emerg Infect Dis. 17(7), 1209–1215. DOI: 10.3201/eid1707.101509.

LeibundGut-Landmann, S., Weidner, K., Hilbi, H., and Oxenius, A. (2011). Nonhematopoietic cells are key players in innate control of bacterial airway infection. J Immunol. 186(5), 3130–3137. Published online 2011/01/29 DOI: 10.4049/jimmunol.1003565.

Lück, P.C., Freier, T., Steudel, C., Knirel, Y.A., Lüneberg, E., Zähringer, U., and Helbig, J.H. (2001). A point mutation in the active site of *Legionella pneumophila O*-acetyltransferase results in modified lipopolysaccharide but does not influence virulence. Int J Med Microbiol. 291(5), 345–352. DOI: 10.1078/1438-4221-00140.

Lück, P.C., Schuppler, M., and Helbig, J.H. (2002). Changes in the lag-1 locus of *Legionella pneumophila* serogroup 1 strains result in different lipopolysaccharides recognized by monoclonal antibodies but do not influence virulence. In: Legionella, (American Society of Microbiology), pp. 52–55.

Luneberg, E., Mayer, B., Daryab, N., Kooistra, O., Zahringer, U., Rohde, M., Swanson, J., and Frosch, M. (2001). Chromosomal insertion and excision of a 30 kb unstable genetic element is responsible for phase variation of lipopolysaccharide and other virulence determinants in *Legionella pneumophila*. Mol Microbiol. 39(5), 1259–1271. Published online 2001/03/17 DOI: 10.1111/j.1365-2958.2001.02314.x.

Lüneberg, E., Zähringer, U., Knirel, Y.A., Steinmann, D., Hartmann, M., Steinmetz, I., Rohde, M., Köhl, J., and Frosch, M. (1998). Phase-variable expression of lipopolysaccharide contributes to the virulence of *Legionella pneumophila*. J Exp Med. 188(1), 49–60.

Macfarlane, J.T., and Worboys, M. (2012). Showers, sweating and suing: Legionnaires’ disease and ‘new’ infections in Britain, 1977–90. Med Hist. 56(1), 72–93. DOI: 10.1017/S0025727300000284.

Mascarenhas, D.P., Pereira, M.S., Manin, G.Z., Hori, J.I., and Zamboni, D.S. (2015). Interleukin 1 receptor-driven neutrophil recruitment accounts to MyD88-dependent pulmonary clearance of *Legionella pneumophila* infection *in vivo*. J Infect Dis. 211(2), 322–330. Published online 2014/08/12 DOI: 10.1093/infdis/jiu430.

Mintz, C.S., Arnold, P.I., Johnson, W., and Schultz, D.R. (1995). Antibody-independent binding of complement component C1q by *Legionella pneumophila*. Infect Immun. 63(12), 4939–4943.

Mintz, C.S., Schultz, D.R., Arnold, P.I., and Johnson, W. (1992). *Legionella pneumophila* lipopolysaccharide activates the classical complement pathway. Infect Immun. 60(7), 2769–2776.

Moller-Kristensen, M., Ip, W.K., Shi, L., Gowda, L.D., Hamblin, M.R., Thiel, S., Jensenius, J.C., Ezekowitz, R.A., and Takahashi, K. (2006). Deficiency of mannose-binding lectin greatly increases susceptibility to postburn infection with *Pseudomonas aeruginosa*. J Immunol. 176(3), 1769–1775.

N. Whitfield, N., and S. Swanson, M. (2006). *Lag-1* Acetylation of Lipopolysaccharide. In: Legionella (American Society of Microbiology).

Newton, H.J., Ang, D.K., van Driel, I.R., and Hartland, E.L. (2010). Molecular pathogenesis of infections caused by *Legionella pneumophila*. Clin Microbiol Rev. 23(2), 274–298. DOI: 10.1128/CMR.00052-09.

O’Connor, T.J., Adepoju, Y., Boyd, D., and Isberg, R.R. (2011). Minimization of the *Legionella pneumophila* genome reveals chromosomal regions involved in host range expansion. Proc Natl Acad Sci U S A. 108(36), 14733–14740. Published online 2011/08/30 DOI: 10.1073/pnas.1111678108.

Page, A.J., Cummins, C.A., Hunt, M., Wong, V.K., Reuter, S., Holden, M.T., Fookes, M., Falush, D., Keane, J.A., and Parkhill, J. (2015). Roary: rapid large-scale prokaryote pan genome analysis. Bioinformatics. 31(22), 3691–3693. Published online 2015/07/20 DOI: 10.1093/bioinformatics/btv421.

Parr, A., Whitney, E.A., and Berkelman, R.L. (2015). Legionellosis on the Rise: A Review of Guidelines for Prevention in the United States. J Public Health Manag Pract. 21(5), E17–26. DOI: 10.1097/PHH.0000000000000123.

Petzold, M., Thürmer, A., Menzel, S., Mouton, J.W., Heuner, K., and Lück, C. (2013). A structural comparison of lipopolysaccharide biosynthesis loci of *Legionella pneumophila* serogroup 1 strains. BMC Microbiol. 13, 198. Published online 2013/09/04 DOI: 10.1186/1471-2180-13-198.

Plouffe, J.F., Para, M.F., and Fuller, K.A. (1985). Serum bactericidal activity against *Legionella pneumophila*. J Clin Microbiol. 22(5), 863–864.

Qin, T., Zhang, W., Liu, W., Zhou, H., Ren, H., Shao, Z., Lan, R., and Xu, J. (2016). Population structure and minimum core genome typing of *Legionella pneumophila*. Sci Rep. 6, 21356. Published online 2016/02/18 DOI: 10.1038/srep21356.

Schrammel, B., Cervero-Aragó, S., Dietersdorfer, E., Walochnik, J., Lück, C., Sommer, R., and Kirschner, A. (2018). Differential development of Legionella sub-populations during short- and long-term starvation. Water Res. 141, 417–427. Published online 2018/04/11 DOI: 10.1016/j.watres.2018.04.027.

Skippington, E., and Ragan, M.A. (2012). Phylogeny rather than ecology or lifestyle biases the construction of *Escherichia coli*-*Shigella* genetic exchange communities. Open Biol. 2(9), 120112. DOI: 10.1098/rsob.120112.

Strassmann, J.E., and Shu, L. (2017). Ancient bacteria-amoeba relationships and pathogenic animal bacteria. PLoS Biol. 15(5), e2002460. Published online 2017/05/02 DOI: 10.1371/journal.pbio.2002460.

Tateda, K., Moore, T.A., Deng, J.C., Newstead, M.W., Zeng, X., Matsukawa, A., Swanson, M.S., Yamaguchi, K., and Standiford, T.J. (2001a). Early recruitment of neutrophils determines subsequent T1/T2 host responses in a murine model of *Legionella pneumophila* pneumonia. J Immunol. 166(5), 3355–3361. Published online 2001/02/24 DOI: 10.4049/jimmunol.166.5.3355.

Tateda, K., Moore, T.A., Newstead, M.W., Tsai, W.C., Zeng, X., Deng, J.C., Chen, G., Reddy, R., Yamaguchi, K., and Standiford, T.J. (2001b). Chemokine-dependent neutrophil recruitment in a murine model of *Legionella pneumonia*: potential role of neutrophils as immunoregulatory cells. Infect Immun. 69(4), 2017–2024. Published online 2001/03/20 DOI: 10.1128/IAI.69.4.2017-2024.2001.

Underwood, A.P., Jones, G., Mentasti, M., Fry, N.K., and Harrison, T.G. (2013). Comparison of the *Legionella pneumophila* population structure as determined by sequence-based typing and whole genome sequencing. BMC Microbiol. 13, 302. Published online 2013/12/24 DOI: 10.1186/1471-2180-13-302.

Verbrugh, H.A., Lee, D.A., Elliott, G.R., Keane, W.F., Hoidal, J.R., and Peterson, P.K. (1985). Opsonization of *Legionella pneumophila* in human serum: key roles for specific antibodies and the classical complement pathway. Immunology. 54(4), 643–653. Published online 1985/04/01.

Watford, W.T., Ghio, A.J., and Wright, J.R. (2000). Complement-mediated host defense in the lung. Am J Physiol Lung Cell Mol Physiol. 279(5), L790–798. DOI: 10.1152/ajplung.2000.279.5.L790.

Wolter, N., Carrim, M., Cohen, C., Tempia, S., Walaza, S., Sahr, P., de Gouveia, L., Treurnicht, F., Hellferscee, O., Cohen, A.L., et al. (2016). Legionnaires’ Disease in South Africa, 2012–2014. Emerg Infect Dis. 22(1), 131–133. DOI: 10.3201/eid2201.150972.

Yu, V.L., Plouffe, J.F., Pastoris, M.C., Stout, J.E., Schousboe, M., Widmer, A., Summersgill, J., File, T., Heath, C.M., Paterson, D.L., et al. (2002). Distribution of *Legionella* species and serogroups isolated by culture in patients with sporadic community-acquired legionellosis: an international collaborative survey. J Infect Dis. 186(1), 127–128. Published online 2002/05/21 DOI: 10.1086/341087.

Ziltener, P., Reinheckel, T., and Oxenius, A. (2016). Neutrophil and Alveolar Macrophage-Mediated Innate Immune Control of *Legionella pneumophila* Lung Infection via TNF and ROS. PLoS Pathog. 12(4), e1005591. Published online 2016/04/23 DOI: 10.1371/journal.ppat.1005591.

