## Supplemental material for "Population analysis of *Legionella pneumophila* reveals the basis of resistance to complement-mediated killing"

Supplementary Information Text

**Supplementary Materials and Methods**

**Bacterial culture.** *L. pneumophila* Sg-1 isolates from the SHLMPRL, SMiRL, Glasgow collection were grown on buffered charcoal yeast extract plates (Legionella CYE agar base - Oxoid, BCYE Growth Supplement SR0110 - Oxoid) for 3 days at 37°C. Three colonies were inoculated into 5 ml of supplemented yeast extract broth (YEB, Yeast extract – Fluka, BCYE Growth Supplement SR0110 - Oxoid) for 24 hours at 37°C 200 rpm. Strains were subcultured by transferring 150 µL of the previous inoculation into 15 mL of YEB at 37°C 200 rpm until it reaches O.D600nm ≈ 0.8. When necessary, media was supplemented with 10 µg/mL of chloramphenicol (Sigma Aldrich) for plasmid maintenance. *Escherichia coli* cells were cultured at 37°C in Luria-Bertani (LB) broth or agar (Melford) supplemented with 10 µg/mL of chloramphenicol (Sigma Aldrich) or 100 µg/mL of ampicillin (Sigma Aldrich). For blue-white colony screening, LB agar plates were spread with 2% X-gal (Melford).

**Whole genome sequencing.** 474 *Legionella* isolates from the *Scottish Haemophilus, Legionella, Meningococcus* & *Pneumococcus* Reference Laboratory (SHLMPRL) isolate collection were recovered from frozen stocks and sent to MicrobesNG (microbesng.uk, Birmingham, United Kingdom) for DNA extraction and whole genome sequencing as follows: plated cultures of each isolate were inoculated into a suspension of plastic beads in a cryopreservative (Microbank™, Pro-Lab Diagnostics UK, United Kingdom). Three beads were washed with extraction buffer containing lysozyme (final concentration 0.1 mg/mL) and RNase A (ITW Reagents, Barcelona, Spain) (final concentration 0.1 mg/mL), incubated for 25 min at 37°C. Proteinase K (VWR Chemicals, Ohio, USA) (final concentration 0.1 mg/mL) and SDS (Sigma-Aldrich, Missouri, USA) (final concentration 0.5% v/v) were added and incubated for 5 min at 65°C. Genomic DNA was purified using an equal volume of SPRI beads and resuspended in EB buffer (Qiagen, Germany).

DNA was quantified with the Quant-iT dsDNA HS kit (ThermoFisher Scientific) assay in an Eppendorf AF2200 plate reader (Eppendorf UK Ltd, United Kingdom). Genomic DNA libraries were prepared using the Nextera XT Library Prep Kit (Illumina, San Diego, USA) following the manufacturer’s protocol with the following modifications: 2 ng of DNA were used as input, and PCR elongation time was increased to 1 min from 30 s. DNA quantification and library preparation were carried out on a Hamilton Microlab STAR automated liquid handling system (Hamilton Bonaduz AG, Switzerland). Pooled libraries were quantified using the Kapa Biosystems Library Quantification Kit for Illumina on a Roche light cycler 96 qPCR machine. Libraries were sequenced with the Illumina HiSeq using a 250 bp paired end protocol.

**Sequence processing and analysis.** Reads were trimmed with Trimmomatic using default settings and *de novo* assembly was performed using SPAdes (v3.10.0) and contigs were reordered with Mauve Contig Reorderer (Rissman *et al.*, 2009). Gene annotation and functional prediction for each assembly was generated using Prokka (v1.12). The quality of whole genome sequencing was assessed using Quast (v4.5) and genomes that exceeded the following metrics were selected as high-quality assemblies for analyses. i) Minimum 1.5 Mbp aligned to the *L. pneumophila* Philadelphia reference (total aligned length), ii) Maximum duplication ratio of 1.03 (based on distribution), iii) Minimum total length of contigs larger than 50 Kbp is 1 Mbp. To maximise the fraction of core genome that can be aligned, isolates belonging to the minor subspecies, (*L. pneumophila* subsp. *fraseri*, *L. pneumophila* subspecies *pascullei* and *L. pneumophila* subspecies *raphaeli*) were not included in downstream analyses. Whole genome sequences are available on the European Nucleotide Archive (ENA) database under Bioproject number PRJEB31628. Functional prediction of non-synonymous mutations in *lag-1* were predicted using PROVEAN (Choi *et al.*, 2012).

12 genes from the LPS cluster, specific to Sg-1 and overlapped by k-mers enriched in clinical isolates (*lpg0762*, *lpg0766*, *lpg0767*, *lpg0768*, *lpg0771*, *lpg0772*, *lpg0773*, *lpg0777*, *lpg0778*, *lpg0779*, *lpg0780*, *lpg0781*). Orthologs of these 12 genes were identified. Genes that were not conserved in all 23 genomes analysed were removed (*lpg0771* and *lpg0777*). Paralogs of *lpg0778* were also removed. Genes were translated and aligned at the codon level using TranslatorX and Mafft. Alignments for all 12 genes were concatenated. Phylogenetic networks were generated using uncorrected P-distances with the equal angle method in Splitstree v4.14.6 (Huson and Bryant, 2006).

Sequence alignments were visualised in Artemis Comparison Tool v.17.0.1 and figures generated with Easyfig v2.2.2 (Carver *et al.*, 2005; Sullivan *et al.*, 2011).

**Genome Wide Association Analyses.** 506 high-quality publicly available genome assemblies of *L. pneumophila* subsp. *pneumophila* were downloaded on the 2^nd^ of March 2017 for inclusion in these analyses. Quality criteria for inclusion of published genomes: not more less than 200 contigs of more than 1Kb, less than 1.03 duplication ratio, not a subspecies or taxonomic outlier (bright green or grey clade), contigs >50 Kb add up to at least 1Mbp, not more than 700 uncalled bases per 100 Kb. At least 3.2 Mbp in length.

From the combined dataset of 900 isolates, we were able to extract metadata describing the isolation source for 758 isolates (environmental or clinical). A ML phylogenetic tree was then constructed from the 1.8 Mbp core genome alignment generated using Parsnp v1.2 (https://github.com/marbl/parsnp). This phylogenetic tree was used as input for the subsampling approach. An automated subsampling algorithm was used to reduce the redundancy of the dataset by iteratively removing one of each pair of isolates which had the same source type. We implemented this algorithm in a modified version of a phylogeny-based dataset reduction tool called Treemmer (Menardo *et al.*, 2018). The modified subsampling script is available on github.com/bawee/Treemmer. Specifically, we added the ability to take into account phenotypic categories (i.e. clinical or environmental) before iteratively removing one taxon from each pair of the most closely related taxa in the phylogeny. This was repeated until the minimum distance between the pairs of isolates with the same phenotype reached a user-defined threshold. This approach was performed with increasing minimum patristic distance thresholds between pairs on the tree, ranging from 0.0001 (≈180 SNPs), 0.001 (≈1800) and 0.01 (≈18000 SNPs). The subsampled datasets contained 452 (60%), 382 (50%) and 353 (47%) of the original isolates, respectively.

GWAS analysis based on *k*-mer enrichment was performed using SEER (Lees *et al.*, 2016). The frequency of *k-*mers between 12 and 100 bp long were counted with fsm-lite (-m 12 -M 100). Only *k*-mers present in 5% and 95% of the total number of isolates (-s <5%> -S <95%>) were used and -maf 0.05 (minimum allele frequency) was used when performing the SEER analysis. The frequency and distribution in clinical and environmental isolates were evaluated individually for each *k-*mer. Pan-GWAS was performed using the homolog clusters generated by ROARY using the following settings (-i 95 -s) (Page *et al.*, 2015). The association analyses were calculated using SCOARY (Brynildsrud *et al.*, 2016). The following thresholds were used to identify the highest-scoring associations: Naïve p-value <0.05, p-value after Bonferroni correction <0.05, Benjamini-Hochberg p-value <0.05 and Empirical p-value after 500 label-switching permutations <0.05 (-e 500).

**Molecular Biology.** For *lag-1* cloning, primers (Variant 1: Fwd- GGC C**GA** **ATT** **C**GT AAG GAA AAA TAA TTT ATC, Rev- ccc **gga tcc** tta tgt tga ata agc taa ctt gtt tga tgt; Variant 2: Fwd – GGC C**GA ATT C**GT AAG GAA AAA TAA TTT ATC, Rev – AT**G GAT CC**T TAC ATC ATC ACC ATC ATC ATT GTT GAA TAA GCC AAC; Variant 3: Fwd – GGC C**GA ATT C**AC ATG CAA GAA TAA TTTA, Rev – ccc **gga tcc** gct tac gta ata taa gct aac tta ttt gat gtg*Eco*RI and *Bam*HI restriction sites in bold) were used to amplify a 1221 bp fragment of chromosomal DNA from *L. pneumophila* strains 4454, 4449 and 4471, encoding the variant 1, 2 and 3 of *lag-1* allele and upstream region, respectively. These fragments were blunt cloned with StrataClone Blunt PCR Cloning kit (Agilent) into vector pSC-B and transformed into *E. coli*. Plasmids from transformants were purified with Monarch Plasmid DNA Miniprep Kit and digested with *Eco*RI and *Bam*HI enzymes (New England BioLabs). *lag-1* gene was ligated to pMMB207 (kind gift from Carmen Buchrieser)using T4 DNA ligase (New England BioLabs) and both pMMB207::*lag-1* and empty plasmid were electroporated into the electrocompetent mAb 3/1-negative strains 4681 and 4312 using a Gene Pulser II (Bio-Rad) following published protocols (Chen *et al.*, 2006). To confirm *O*-acetyltransferase activity of the *lag-1* cloned bacteria, transformed and positive control strains were fixed in 10% formalin (VWR) for 20 min and incubated with 1:100 of mouse mAb 3/1 (Dresden panel) for 30 min in ice prior incubation with 1:100 polyclonal AlexaFluor 488 conjugated donkey anti-mouse IgG (H+L) (ThermoFisher Scientific). After incubation on ice for 30 min, the bacteria were washed and resuspended in PBS for flow cytometry analysis. DsRed fluorescen protein expression was introduced into WT *lag‑1* positive and negative strains by electroporation with pSW001 plasmid (pMMB207C, ΔlacIq, constitutive *dsred*, kind gift from Carmen Buchrieser) (Rolando *et al.*, 2013). DsRed expression was confirmed by flow cytometry. Samples were analysed on a BD LSRFortessa X-20 flow cytometer (BD biosciences) and data analysed using FlowJo software (v10).

**Ethics statement.** Ethical approval for the collection of blood from anonymous donors was granted by the University of Edinburgh Research Ethics Committee. This study was reviewed by the University of Edinburgh, College of Medicine Ethics Committee (2009/01) and subsequently renewed by the Lothian Research Ethics Committee (11/AL/0168). Written informed consent was received from all volunteers participating in the study.

**Plasma and serum killing assays.** Human blood was obtained from healthy volunteers in syringes treated with anticoagulant citrate dextrose. Plasma was obtained by centrifuging whole blood for 15 min at 25°C at 1200g without the centrifuge brake and collecting the layer above the buffy coat. Normal serum was collected from human blood obtained from healthy volunteers in BD Vacutainer serum tubes. Tubes were centrifuged for 10 min at 25°C 1200g and the supernatant collected. Plasma and serum were stored following immediate freezing at -80°C. Samples used were negative for anti-*L. pneumophila* antibodies as determined by standard diagnostic serology methods (immune fluorescence test). The ability of *L. pneumophila* strains to resist killing by human plasma and serum was determined by incubating 1x10^5^ bacterial cells in 90% plasma or serum at 37°C for 60 min. Serial dilutions were plated in duplicate on BCYE media, incubated at 37°C for 3 days before enumeration. Heat inactivation of plasma was carried out by incubation at 56°C for 30 min in a water bath. Both plasma, serum, heat-inactivated plasma and heat-inactivated serum were centrifuged at 4000g for 20 min at 4°C prior use. Inhibition of complement pathways was carried out by adding 12.5 mM EDTA (Sigma Aldrich), 12.5 mM EGTA/Mg^2+^ (Sigma Aldrich) or 100 mM mannose (Acros Organics) to normal human serum or by using C1q- and Factor B- depleted serum (Pathway Diagnostics).

**C3 binding assays.** For analysis of C3 deposition on the surface of *L. pneumophila*, bacteria were grown to post-exponential phase and fixed in 10% formalin (VWR) for 20 min. Nunc maxisorp ELISA plates were coated with 1x10^7^ bacteria per well overnight at 4°C, blocked with 1% w/v bovine serum albumin (Sigma Aldrich) in PBS for 2 hours and incubated with serial dilution of human serum for 2 hours at room temperature. Binding of C3 at the surface of the bacteria was determined by incubating with 1:100 FITC conjugated goat F(ab’)2 anti-human C3 complement (Protos Immunoresearch) for 2 hours at room temperature and fluorescence detected in using the CLARIOstar fluorescence plate reader (BMG Labtech).

**Bovine Bronchoalveolar Lavage.** *Ex vivo* cow lungs were washed with PBS at room temperature (ref, Lindert). 500 ml of fresh BAL were centrifuged at 500g for 7 min and supernant was filtered through a 0.2 µm pore size membrane filters (Nalgene) to remove any bovine cells or contaminants. BAL proteins were then concentrated on an Amicon centrifugal filters with a 15 kb cut-off to a final volume of 25 ml. Heat inactivation of BAL was carried out by incubation at 56°C for 30 min in a water bath. The ability of *L. pneumophila* strains to resist killing by bovine BAL was determined by incubating 1x10^6^ bacterial cells per mL in concentrated bovine BAL at 37°C for 60 min. Serial dilutions were plated in duplicate on BCYE media, incubated at 37°C for 3 d before enumeration.

**Neutrophil assay.** Human venous blood was obtained from healthy volunteers in syringes treated with anticoagulant citrate dextrose and the isolation of neutrophils was performed by Ficoll/Histopaque centrifugation (Surewaard *et al.*, 2013). *L. pneumophila*-DsRed strains were incubated with 10% normal human serum for 15 min at 37°C before incubation with 2.5x10^5^ purified neutrophils (MOI 10). Infection was maintain at 37°C, 5% CO_2_ with orbital shaking at 600 rpm for 30 min. Cells were then fixed in 10% formalin (VWR) for 20 min and kept in PBS for flow cytometry analysis. Samples were analysed on a BD LSRFortessa X-20 flow cytometer (BD biosciences) and data analysed using FlowJo software (v10).

Fig. S1

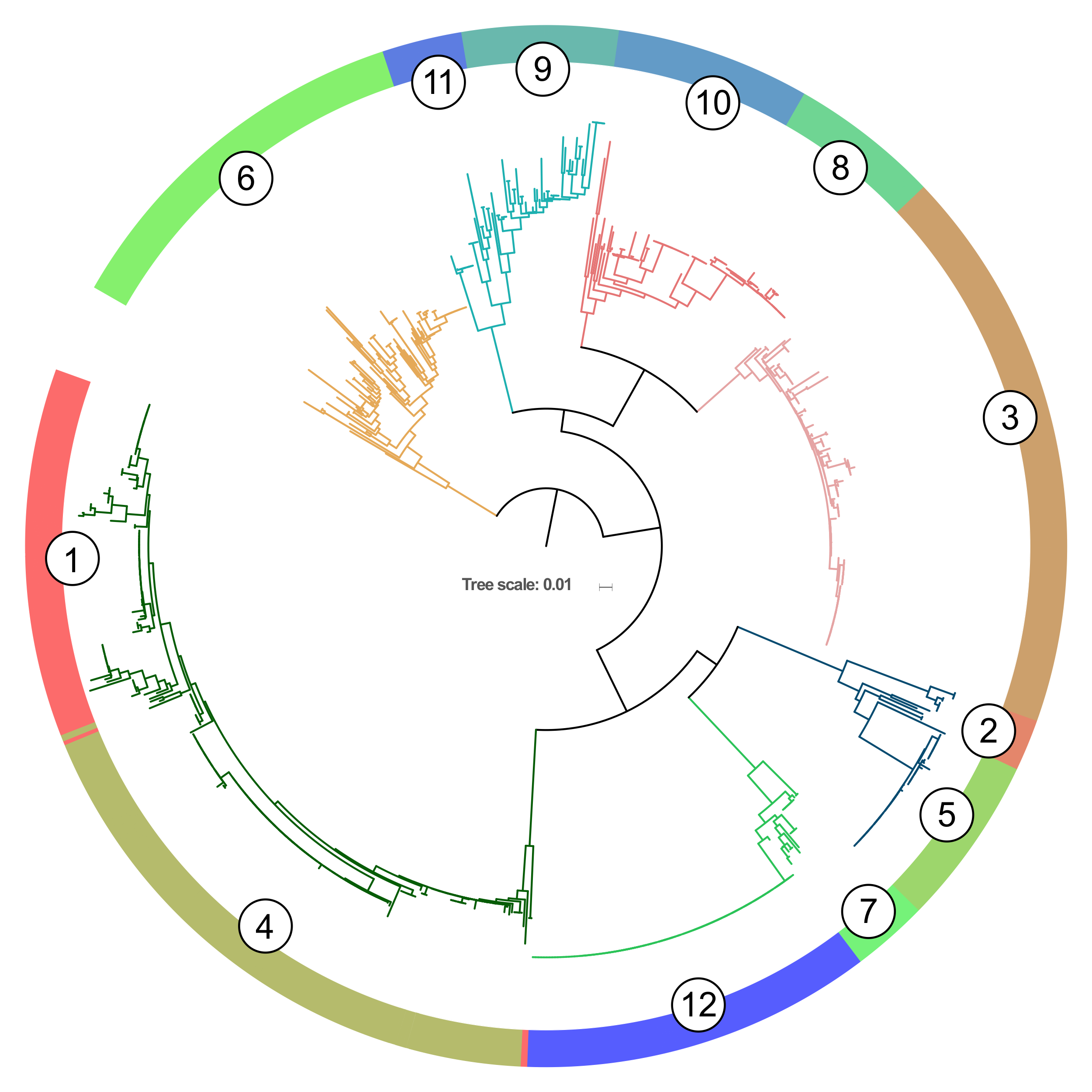

Fig. S1. Maximum likelihood phylogenetic tree with BAPS clusters indicated. Each of the 7 major Maximum-Likelihood phylogenetic clades (branches coloured corresponding to Figure 1) that define the *L. pneumophila* subsp. *pneumophila* population structure is supported by at least 1 and a maximum of 2 Bayesian Analysis of Population Structure (BAPS) clusters determined from the core SNP alignment.

Fig. S2

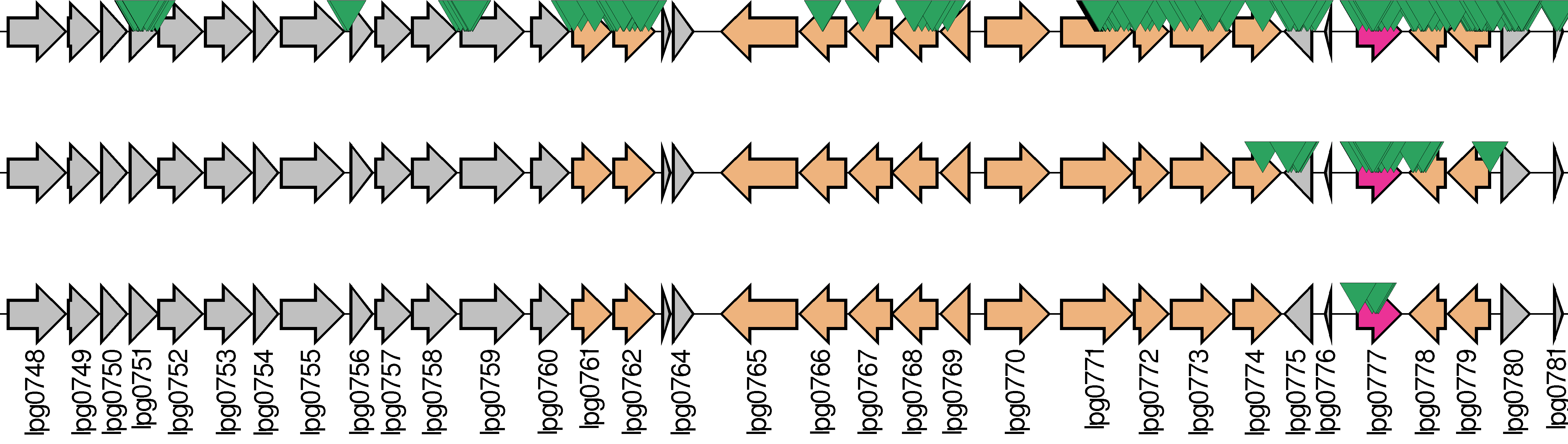

Fig. S2. Localisation of significantly over-represented *k*-mers (indicated by green arrows) detected with SEER to the lipopolysaccharide (LPS) gene cluster (*lpg0748*-*lpg0781*) relative to the Philadelphia 1 reference genome. From top to bottom, the effect of subsampling using increasingly stringent minimum phylogenetic distance thresholds (Top: 0.0001 (≈180 SNPs), Middle: 0.001 (≈1800) and Bottom: 0.01 (≈18000 SNPs).

Fig. S3

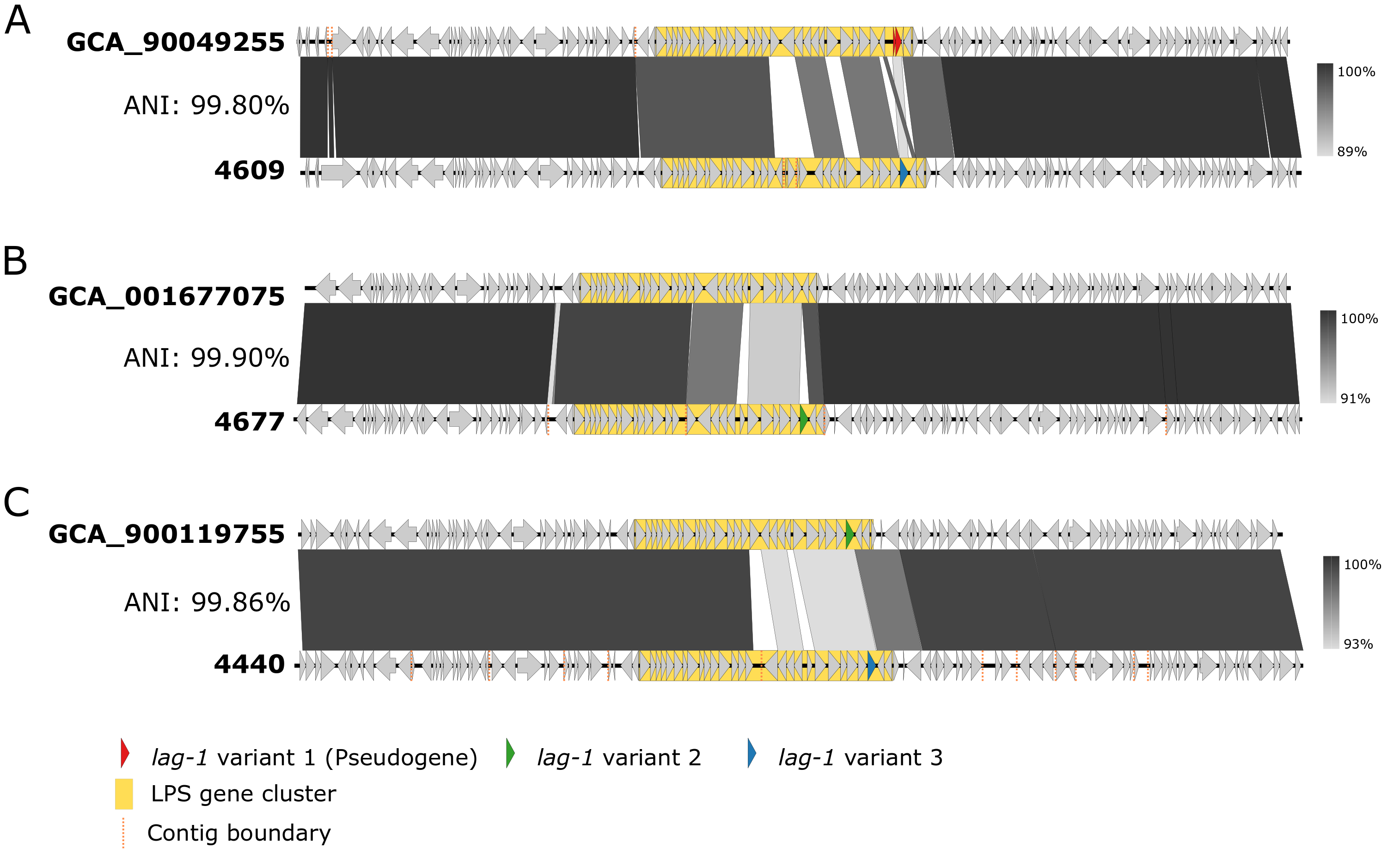

Fig. S3. Recombination events driving replacement of *lag-1* variants. Pairwise alignment of genomic region flanking the LPS gene cluster indicating homologous recombination events between closely related genomes associated with switching between a disrupted *lag-1* variant 1 and a *lag-1* variant 3 (A), a *lag-1* LPS cluster with *lag-1* variant 2 (B), and a *lag-1* variant 2 with a *lag-1* variant 3 (C). The average nucleotide identity (ANI) between genomes is shown on the left of the alignment and the level of nucleotide identity of the region shown is indicated by the grey scale on the right of each alignment.

Fig. S4

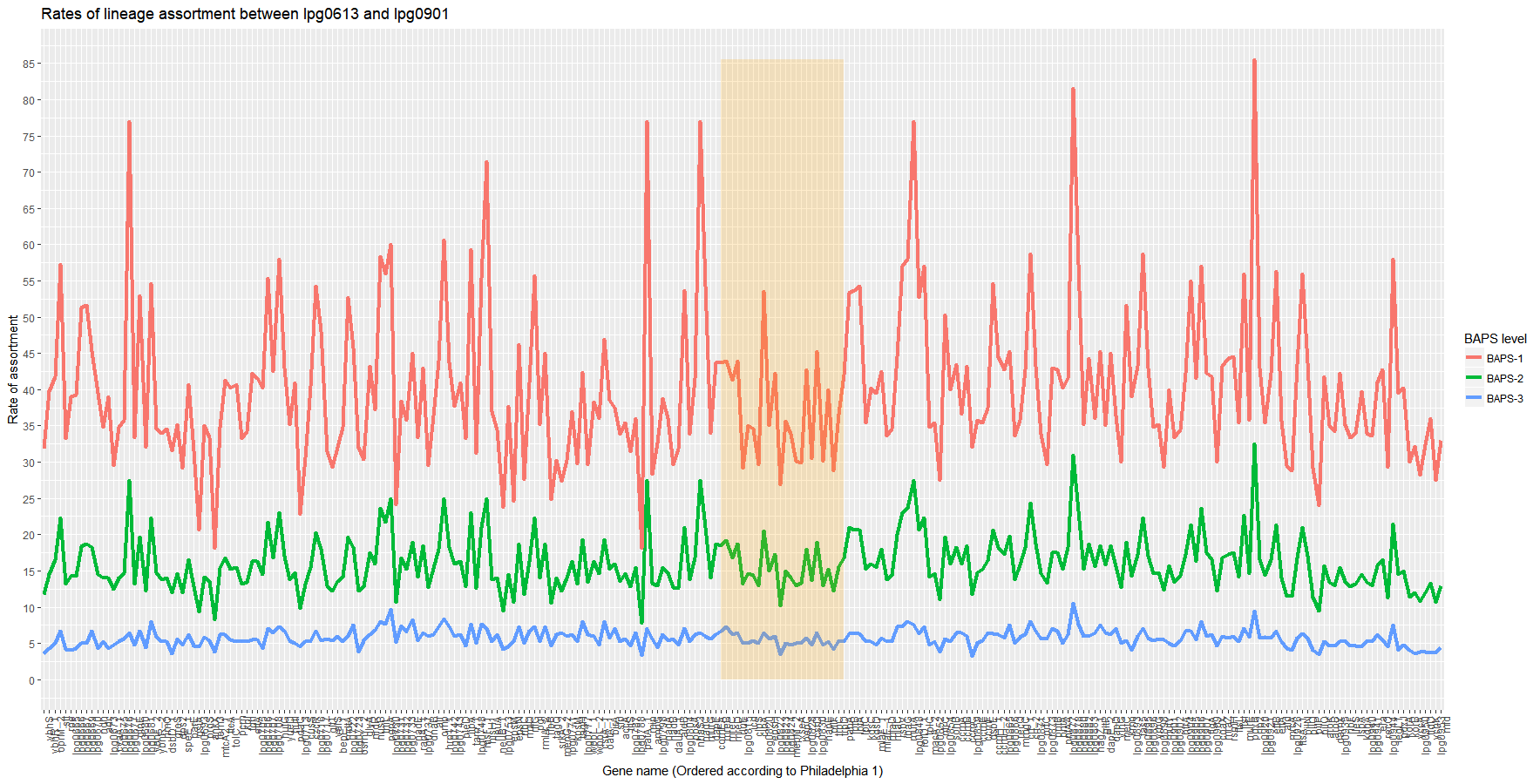

Fig. S4. Rates of assortment across LPS biosynthesis clusters (highlighted) relative to surrounding genes. The rate of assortment is calculated by comparing the number of unique combinations between alleles, as defined by fastGEAR, and the BAPS populations clusters, normalised by the number of possible alleles for each gene.

Fig. S5

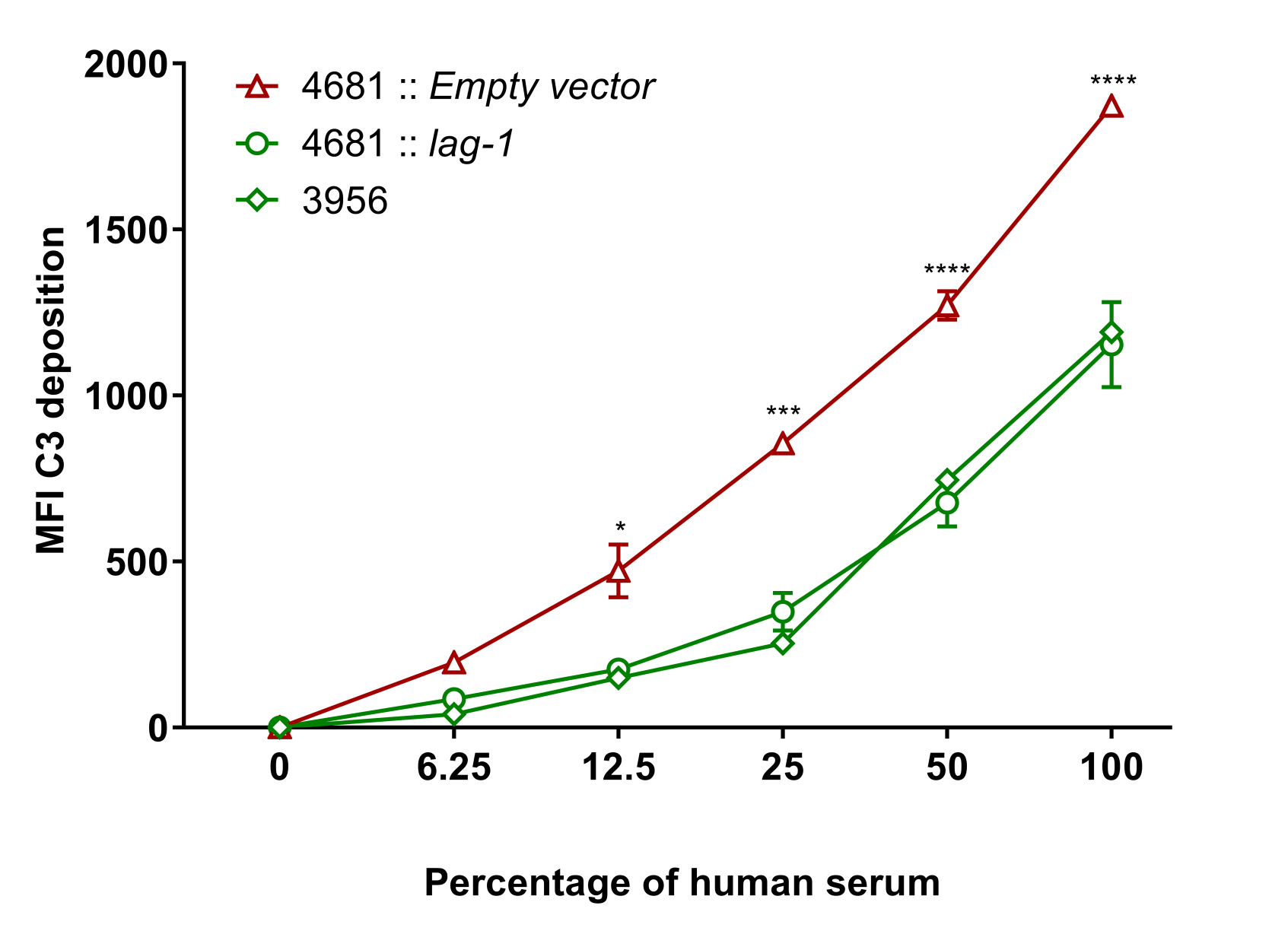

Fig S5. Deposition of human C3 on the surface of *L. pneumophila* is dependent on *lag-1* expression. Fixed bacteria were incubated with serial dilutions of normal human serum for 2 hours prior incubation with FITC conjugated anti-C3 Fab antibodies. Two-way ANOVA multiple comparison test, *p<0.05, ***p<0.001, ****p<0.0001.

**Fig. S6**

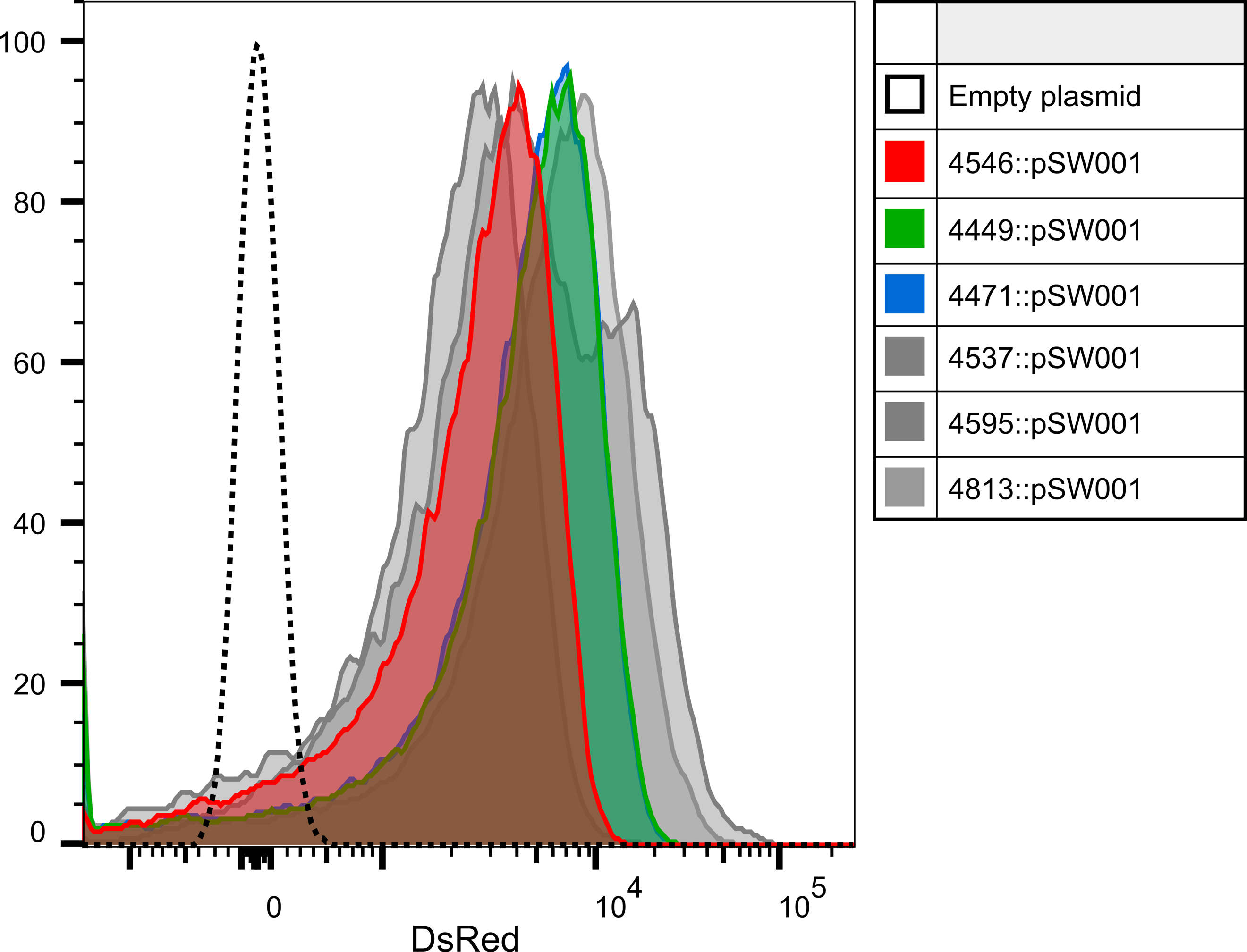

Fig. S6. *L. pneumophila* strains transformed with pSW001 plasmid express similar levels of fluorescent DsRed. DsRed MFI of *L. pneumophila* strains that express *lag-1* variant 1 (red), variant 2 (green), variant 3 (blue) or are negative for *lag-1* expression (grey).

**Table S1. Strain list and metadata for genomes employed in the study**

**(see attached dataset)**

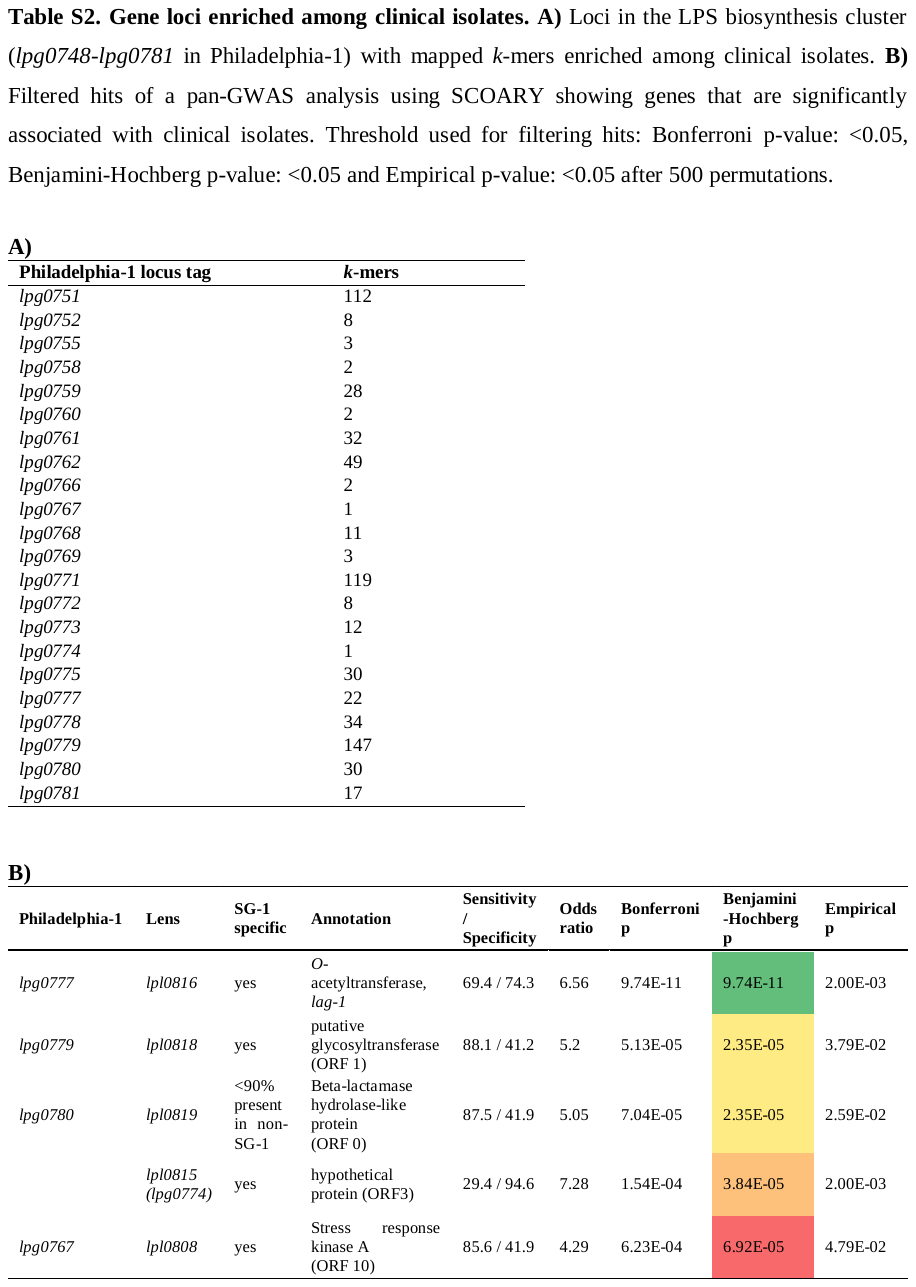

**Supplemental References**

Brynildsrud, O., Bohlin, J., Scheffer, L., and Eldholm, V. (2016). Rapid scoring of genes in microbial pan-genome-wide association studies with Scoary. Genome Biol. 17(1), 238. Published online 2016/11/25 DOI: 10.1186/s13059-016-1108-8.

Carver, T.J., Rutherford, K.M., Berriman, M., Rajandream, M.A., Barrell, B.G., and Parkhill, J. (2005). ACT: the Artemis Comparison Tool. Bioinformatics. 21(16), 3422-3423. Published online 2005/06/25 DOI: 10.1093/bioinformatics/bti553.

Chen, D.Q., Huang, S.S., and Lu, Y.J. (2006). Efficient transformation of *Legionella pneumophila* by high-voltage electroporation. Microbiol Res. 161(3), 246-251. DOI: 10.1016/j.micres.2005.09.001.

Choi, Y., Sims, G.E., Murphy, S., Miller, J.R., and Chan, A.P. (2012). Predicting the functional effect of amino acid substitutions and indels. PLoS One. 7(10), e46688. Published online 2012/10/08 DOI: 10.1371/journal.pone.0046688.

Huson, D.H., and Bryant, D. (2006). Application of phylogenetic networks in evolutionary studies. Mol Biol Evol. 23(2), 254-267. DOI: 10.1093/molbev/msj030.

Lees, J.A., Vehkala, M., Välimäki, N., Harris, S.R., Chewapreecha, C., Croucher, N.J., Marttinen, P., Davies, M.R., Steer, A.C., Tong, S.Y., et al. (2016). Sequence element enrichment analysis to determine the genetic basis of bacterial phenotypes. Nat Commun. 7, 12797. Published online 2016/09/16 DOI: 10.1038/ncomms12797.

Menardo, F., Loiseau, C., Brites, D., Coscolla, M., Gygli, S.M., Rutaihwa, L.K., Trauner, A., Beisel, C., Borrell, S., and Gagneux, S. (2018). Treemmer: a tool to reduce large phylogenetic datasets with minimal loss of diversity. BMC Bioinformatics. 19(1), 164. Published online 2018/05/02 DOI: 10.1186/s12859-018-2164-8.

Page, A.J., Cummins, C.A., Hunt, M., Wong, V.K., Reuter, S., Holden, M.T., Fookes, M., Falush, D., Keane, J.A., and Parkhill, J. (2015). Roary: rapid large-scale prokaryote pan genome analysis. Bioinformatics. 31(22), 3691-3693. Published online 2015/07/20 DOI: 10.1093/bioinformatics/btv421.

Rissman, A.I., Mau, B., Biehl, B.S., Darling, A.E., Glasner, J.D., and Perna, N.T. (2009). Reordering contigs of draft genomes using the Mauve aligner. Bioinformatics. 25(16), 2071-2073. Published online 2009/06/12 DOI: 10.1093/bioinformatics/btp356.

Rolando, M., Sanulli, S., Rusniok, C., Gomez-Valero, L., Bertholet, C., Sahr, T., Margueron, R., and Buchrieser, C. (2013). *Legionella pneumophila* effector RomA uniquely modifies host chromatin to repress gene expression and promote intracellular bacterial replication. Cell Host Microbe. 13(4), 395-405. Published online 2013/04/23 DOI: 10.1016/j.chom.2013.03.004.

Sullivan, M.J., Petty, N.K., and Beatson, S.A. (2011). Easyfig: a genome comparison visualizer. Bioinformatics. 27(7), 1009-1010. Published online 2011/02/01 DOI: 10.1093/bioinformatics/btr039.

Surewaard, B.G., van Strijp, J.A., and Nijland, R. (2013). Studying interactions of Staphylococcus aureus with neutrophils by flow cytometry and time lapse microscopy. J Vis Exp. (77), e50788. Published online 2013/07/31 DOI: 10.3791/50788.

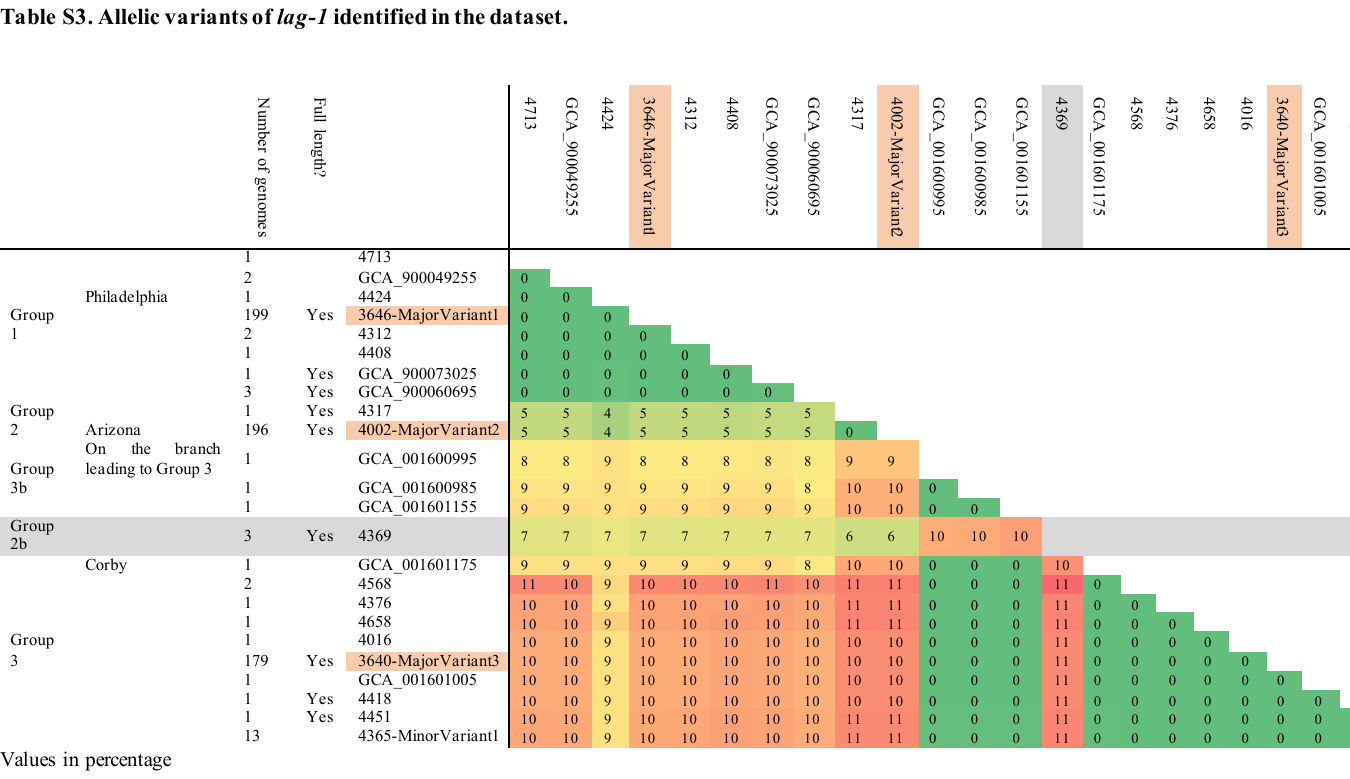

**Table S4. Relative reassortment rates of genes conserved in at least 30% of genomes**

**(see attached dataset)**
